# The Nup2 meiotic-autonomous region relieves autoinhibition of Nup60 to promote progression of meiosis and sporulation in *Saccharomyces cerevisiae*

**DOI:** 10.1101/2021.10.12.463977

**Authors:** Kelly Komachi, Sean M. Burgess

## Abstract

During meiosis, chromosomes undergo dramatic changes in structural organization, nuclear positioning, and motion. Although the nuclear pore complex has been shown to affect genome organization and function in vegetative cells, its role in meiotic chromosome dynamics has remained largely unexplored. Recent work in the budding yeast *Saccharomyces cerevisiae* demonstrated that the mobile nucleoporin Nup2 is required for normal progression through meiosis I prophase and sporulation in strains where telomere-led chromosome movement has been compromised. The meiotic autonomous region (MAR), a short fragment of Nup2 responsible for its role in meiosis, was shown to localize to the nuclear envelope via Nup60 and to bind to meiotic chromosomes. To understand the relative contribution these two activities have on MAR function, we first carried out a screen for MAR mutants defective in sporulation and found that all the mutations disrupt interaction with both Nup60 and meiotic chromosomes. Moreover, *nup60* mutants phenocopy *nup2* mutants, exhibiting similar nuclear division kinetics, sporulation efficiencies, and genetic interactions with mutations that affect the telomere bouquet. Although full-length Nup60 requires Nup2 for function, removal of Nup60’s C-terminus allows Nup60 to bind meiotic chromosomes and promote sporulation without Nup2. In contrast, binding of the MAR to meiotic chromosomes is completely dependent on Nup60. Our findings uncover an inhibitory function for the Nup60 C-terminus and suggest that Nup60 mediates recruitment of meiotic chromosomes to the nuclear envelope, while Nup2 plays a secondary role counteracting Nup60’s autoinhibition.

## Introduction

The chromosome events of meiosis, including pairing, synapsis, and crossing over between homologous chromosomes, take place in a crowded nuclear environment. It is not well understood if or how the physical location of sequences in the nucleus contribute to meiotic processes. One organizing feature of chromosomes specific to meiosis is the telomere bouquet, in which the chromosome ends cluster at the nuclear envelope by attaching to the linker of nucleoskeleton and cytoskeleton (LINC) complex (Burke 2018). The indirect attachment of telomeres to cytoskeletal motor proteins results in dramatic chromosome movements that coincide with the process of homolog pairing. Whether other chromosome regions associate with the nuclear envelope is unknown.

In budding yeast, the Ndj1 protein acts as an adapter between the telomeres and the LINC complex. Mutations in *NDJ1* disrupt the telomere bouquet, abolish telomere-led chromosome movements, and lead to defects in pairing (Scherthan *et al*. 2007; Conrad *et al*. 2008; Wanat *et al*. 2008). Nevertheless, *ndj1* mutants have nearly wild-type sporulation efficiency and spore viability, implying that yeast might have an auxiliary system that compensates for loss of the telomere bouquet. The nucleoporin Nup2 emerged as a likely component of this potential backup mechanism from studies that uncovered a synthetic interaction between *nup2* and *ndj1* mutations, suggesting that the LINC complex and the nuclear pore complex (NPC) promote meiosis and sporulation by two functionally redundant pathways (Chu *et al*. 2017).

Nup2 is part of the nuclear basket, a fibrous structure on the nucleoplasmic face of the NPC that also contains the proteins Nup60, Nup1, Mlp1, and Mlp2 (Raices and D’Angelo 2021). Although first characterized as a component of the transport machinery, Nup2 was later found to play roles in chromatin organization, gene regulation, and DNA damage repair (Ishii *et al*. 2002; Dilworth *et al*. 2005; Kim *et al*. 2017; Brickner *et al*. 2019). Deletion analysis showed that a short fragment of Nup2 is necessary and sufficient for its meiotic function. This meiotic autonomous region (MAR) of Nup2 binds to both the nuclear envelope and meiotic chromosomes (Chu *et al*. 2017), suggesting that Nup2 might be helping to organize chromosomes by tethering them to the NPC. There are numerous examples in vegetative cells where Nup2 helps bring promoter regions of activated genes to the NPC (Casolari *et al*. 2004; Dilworth *et al*. 2005; Schmid *et al*. 2006; Brickner *et al*. 2012, 2019; Kim *et al*. 2017). In addition, the MAR shares weak homology with a segment of *Aspergillus nidulans* Nup2 that tethers the NPC to chromatin during mitosis (Markossian *et al*. 2015; Suresh *et al*. 2017, 2018). Thus, it seemed plausible that Nup2 could be acting as a bridge between the nuclear envelope and chromatin during meiosis as well.

However, another possibility is that Nup2’s meiotic function does not require association with the NPC. Nup2 localizes to the inner nuclear membrane by binding to Nup60 but can also be found in the nucleoplasm and is more mobile than the other basket nucleoporins (Solsbacher *et al*. 2000; Dilworth *et al*. 2001; Denning *et al*. 2001). Similarly, Nup50, the metazoan homolog of Nup2, associates with the NPC via Nup153, the metazoan Nup60 homolog (Hase and Cordes 2003; Makise *et al*. 2012), but dissociates with relative ease (Rabut *et al*. 2004). Furthermore, Nup50 can bind to and activate genes in the nucleoplasm (Kalverda *et al*. 2010), and when transcription is inhibited, the bulk of Nup50 moves off the NPC and stably re-localizes to nucleoplasmic chromatin in a Nup153-independent manner (Buchwalter *et al*. 2014).

In this study, we began by addressing the question of whether Nup2 promotes meiosis as a component of the NPC or as a diffusible factor in the nucleoplasm. If presence at the NPC is critical for Nup2’s meiotic function, then disruption of the interaction between Nup2 and Nup60 should interfere with sporulation. Conversely, if Nup2 is acting in the nucleoplasm, then Nup2 is likely to function independently of Nup60. Mutagenesis of the MAR revealed that the ability to bind Nup60 is crucial for MAR function and that the MAR depends on Nup60 for localization to both the nuclear envelope and meiotic chromosomes. Somewhat surprisingly, we found that an N-terminal fragment of Nup60 not only supports sporulation but does so in the absence of Nup2. Since full-length Nup60 does not function in sporulation without Nup2, we conclude that the C-terminus of Nup60 inhibits the activity of its N-terminus and Nup2 relieves that inhibition. Overall, our work suggests that Nup2 and Nup60 work in concert, with Nup60 binding both the nuclear envelope and meiotic chromosomes and with Nup2 acting mainly to prevent Nup60 from inhibiting itself. We propose that the NPC plays a role in sporulation by recruiting meiotic chromosomes to the nuclear envelope and provides a second level of chromosome organization that complements telomere attachment to the nuclear periphery.

## Methods

### Strains and media

All strains in this study except for the Y2H strain are derivatives of SK1 and are listed in Supplemental information Table S1. Plasmids and primers are listed in Tables S2 and S3. Yeast media were prepared as previously described (Lui and Burgess 2009; Chu *et al*. 2017). Standard techniques were used for yeast manipulation. Gene knockouts and fluorescently tagged constructs were created using tailed PCR-based gene replacement and tagging techniques (Longtine *et al*. 1998; Sheff and Thorn 2004). Gene disruptions were confirmed by PCR, and new alleles were confirmed by both PCR and sequencing. Unmarked deletion alleles of *NUP60* were created by transformation of a *nup60::URA3* strain with PCR-generated DNA fragments. Transformants were selected for on plates containing 5-fluoroorotic acid (Boeke *et al*. 1984). All alleles generated by 5-FOA selection were outcrossed twice before further analysis.

### MAR mutant screen

The *nup2(51-175)-GFP::CaURA3 (MAR-GFP::URA3)* fragment from plasmid pSB470 was used as the template for PCR mutagenesis and contains part of the *NUP2* 5’ UTR plus the start codon (−203 to +3), the coding sequence for Nup2 amino acids 51-175, the 2,441 bp *GFP::URA3* fragment from pKT209 (Sheff and Thorn, 2004) starting 24 bp upstream of the GFP start codon, and part of the *NUP2* 3’ UTR (+2161 to +2482). A PCR fragment from *NUP2* (−203) to the GFP start codon was synthesized using Taq polymerase under standard conditions, which introduced errors at a high frequency due to the low fidelity of the enzyme. A second PCR fragment from 24 bp upstream of the GFP start codon to *NUP2* (+2482) was generated using the high-fidelity polymerase Phusion (New England Biolabs). The two fragments were joined by overlap extension PCR (Horton 1989) with the Phusion polymerase and transformed into SBY6259 (*MATa ho::hisG leu2::hisG ura3::hisG his4::LEU2 nup2(Δ51-175) ndj1::TRP1)*. The mutagenized *MAR-GFP::CaURA3* fragment integrated at the *NUP2* locus, replacing *nup2(Δ51-175)*. After three days, Ura^+^ transformants were patched onto fresh -Ura plates alongside positive and negative controls. The patches were grown for 24 hours, then replica plated onto -Ura plates overlaid with a disc of Whatman 1 paper and onto -His-Ura plates containing a mating lawn of SBY6260 (*MATalpha ho::hisG leu2::hisG ura3(ΔSma-PstI) HIS4::LEU2 nup2::KanMx ndj1::KanMx)*. The patches on the Whatman filter were incubated at 30°C for 24 hours, then screened for GFP fluorescence using an ImageQuant LAS 4000. Candidates that had lost GFP fluorescence were eliminated from further analysis since they were likely to contain nonsense mutations or mutations that inactivate GFP. The -His-Ura plates were incubated at 30°C for two days, then the resulting patches of His^+^ Ura^+^ diploids were replica plated onto rich sporulation plates (1% potassium acetate, 0.1% peptone, 0.1% yeast extract, 0.05% glucose, 2% agar) and incubated at 30°C for 3 days. The screen was performed in this manner rather than by transforming a diploid strain because mitochondrial loss during the transformation step led to a high frequency of false positives in trial screens. Sporulation was detected as dityrosine fluorescence as in (McKee and Kleckner 1997) but using an Alpha Innotech AlphaImager 3400 on the reflective UV setting with a SYBR green filter for detection. Sporulation-defective candidates were streaked for single colonies, patched onto -Ura plates and retested. The MAR coding region was amplified by PCR using Phusion polymerase, and the resulting fragments were commercially sequenced (Quintara Biosciences). For all further analysis, the entire *MAR-GFP::URA3* fragment from each of the mutants was amplified by PCR and transformed into SBY1900 to allow construction of strains in the SBY1903 background.

### Preparation of diploid colonies for sporulation assays and chromosome spreads

Glycerol stocks of haploid parent strains were patched onto YPG plates, allowed to mate for 14 hours at 30°C, then streaked onto YPD and grown for 48 hours at 30°C. Diploid colonies were identified based on colony and cell morphology. For experiments involving *nup60* strains where the nibbled colony phenotype (Zhao *et al*. 2004) precluded morphology-based identification, all strains were prepared by patching haploid parents on YPG as above, then streaking the cells onto SC -His -Ura plates to select for diploids. Wild-type strains from colonies grown on SC -His -Ura plates sporulated as efficiently as those from colonies grown on YPD.

### Quantitative sporulation assay

Cells were induced to enter meiosis in liquid culture according to Lui and Burgess 2009. Samples were collected after 24 hours in sporulation medium (SPM) and sonicated for 3 seconds on setting 3 using a Fisher Sonic Dismembrator 550 fitted with a microtip to disperse cell clumps. Samples were examined under white light using a Zeiss 47 30 11 9901 microscope to assess sporulation. Cells with 2 or more spores were counted as sporulated, while cells with 1 or no spores were counted as unsporulated. At least 200 cells total were counted for each sample. For all strains, the assay was performed in triplicate using cultures grown from independent colonies and repeated at least once, such that n ≥ 6. For each strain, the sporulation efficiency is reported as the mean and standard deviation for data pooled from all assays performed on that strain. Complete sporulation data are reported in Supplemental File S1.

### Yeast two-hybrid assay

The coding sequence of the MAR was inserted into the Gal4 DNA-binding domain vector pGBKT7 (Clontech); the coding sequences for full length Nup60 and Nup60(188-388) were separately inserted into the Gal4 activation domain vector pACT2-2 (Arora *et al*. 2004). Bait and prey plasmids were co-transformed into AH109, a strain in which the HIS3 gene is driven by the *GAL1* promoter. Trp^+^ Leu^+^ transformants were inoculated into liquid SC -Trp -Leu media and grown for 24 hours. The cultures were diluted to an OD_600_ of 1.0, and 5 ul were spotted onto SC -Trp -Leu plates and SC -Trp -Leu -His plates containing 25 mM 3-aminotriazole. The plates were grown for 3 days, then imaged using an ImageQuant LAS 4000. At least three independent transformants were tested for each combination of bait and prey plasmids.

### Microscopy and image analysis

All images of whole cells and of meiotic chromosome spreads were obtained with a Zeiss Axioskop epifluorescence microscope equipped with a 100x oil immersion objective and TRITC, FITC, and DAPI filter sets (Chroma). Images were collected with a Hamamatsu C4742-95 camera using Micro-Manager software (Edelstein *et al*. 2010) and were analyzed using Fiji/ImageJ Version 2.0.0-rc-68/1.52e (Schindelin *et al*. 2012).

### Meiotic chromosome spreads

Cells were induced to enter meiosis using the time course protocol cited described above and were collected after 10 hours in sporulation medium. Spheroplasts were prepared as previously described (Grubb *et al*. 2015), then pelleted and washed twice with MES/sorbitol (0.1 M MES-NaOH, pH 6.4, 1 mM EDTA, 0.5 mM MgCl_2_, 1 M sorbitol). Chromosome spreads were prepared and stained according to (Rockmill 2009), with the following modifications. The volumes of 1X MES and FIX used were 160 ul and 440 ul, respectively. The cell mixture was evenly distributed onto 3 slides, which were incubated in a warm, humid chamber for 40 minutes before being washed with Photo-Flo. Spreads were pre-incubated in (PBS + 5% BSA)/fetal bovine serum prior to incubation with primary antibody and in PBS + 5% BSA prior to incubation with secondary antibody. Preincubations were for 30 minutes at room temperature in a humid chamber. Slides were dried for at least 20 minutes at room temperature and stored at −20°C prior to staining. Antibody-stained slides were mounted in Prolong Glass with NucBlue (Thermofisher P36985) and cured for 24 hours at room temperature before imaging. Spreads were repeated for all strains at least once.

### Quantitation of chromosome spread fluorescence

Pachytene nuclei were selected for imaging based on their DNA staining pattern in the DAPI channel. Fluorescence was quantitated from images of spread nuclei as described in (McCloy *et al*. 2014) for images of whole cells. The region of interest (ROI) was defined by manually drawing a line around the NucBlue-stained region in the DAPI channel image. Area and mean fluorescence were measured in the region of interest in the corresponding FITC channel image as well as in background regions adjacent to the ROI. The process was repeated for TRITC channel images when applicable. The total corrected fluorescence (TCF) was calculated as TCF = 0.001 x [integrated density – (area of selected cell × mean fluorescence of background readings)]. At least 10 cells from each strain were imaged and quantified on at least two separate days (biological duplicates). Fluorescence measurements are reported in Files S2-S5. Graphing and statistical analysis was done using Prism 9 GraphPad Software.

### Meiotic time course protocol

The meiotic time course was performed as described in Lui and Burgess 2009. Samples were removed from sporulation media for analysis at 2-hour intervals, starting at t=0 hours. The samples were sonicated as described above for the quantitative sporulation assay, fixed by the addition of ethanol to a final concentration of 40% by volume, and stored at -20°C until analyzed. Samples were stained by adding an equal volume of 1 μg/mL DAPI. Cells with at least two well-defined DAPI-stained foci were counted as multinucleate. At least 200 cells were analyzed for each sample. Duplicate samples were run in parallel on each day and each time course was repeated at least one time on a separate day.

Time course data are reported in File S6.

### Antibodies

The primary antibodies used were chicken polyclonal antibody to GFP (Novus NB100-1614; RRID:AB_10001164) and rabbit polyclonal antibody to mCherry (Novus NBP2-25157; RRID:AB_2753204). The secondary antibodies used were goat anti-chicken 488 (Thermofisher A11039; RRID:AB_2534096) and goat anti-rabbit 594 (Thermofisher A11012; RRID:AB_2534079). All antibodies were used at a 1:1000 dilution. Each antibody was tested for specificity using strains not expressing GFP or mCherry.

## Results

### Point mutations affect MAR function

Previous work had shown that Nup2 and Ndj1 act in functionally redundant pathways to promote meiosis and sporulation. The segment of Nup2 responsible for its function, the MAR, spans amino acids 51-175 (Figure 1A) and has at least two activities: it localizes to the nuclear envelope via its interaction with Nup60, and it binds to meiotic chromosomes (Chu *et al*. 2017). To help clarify whether these activities are important for MAR function, we carried out a screen for mutations that decreased sporulation with the aim of identifying alleles that were defective for either subcellular localization or chromosome binding.

**Figure 1.**
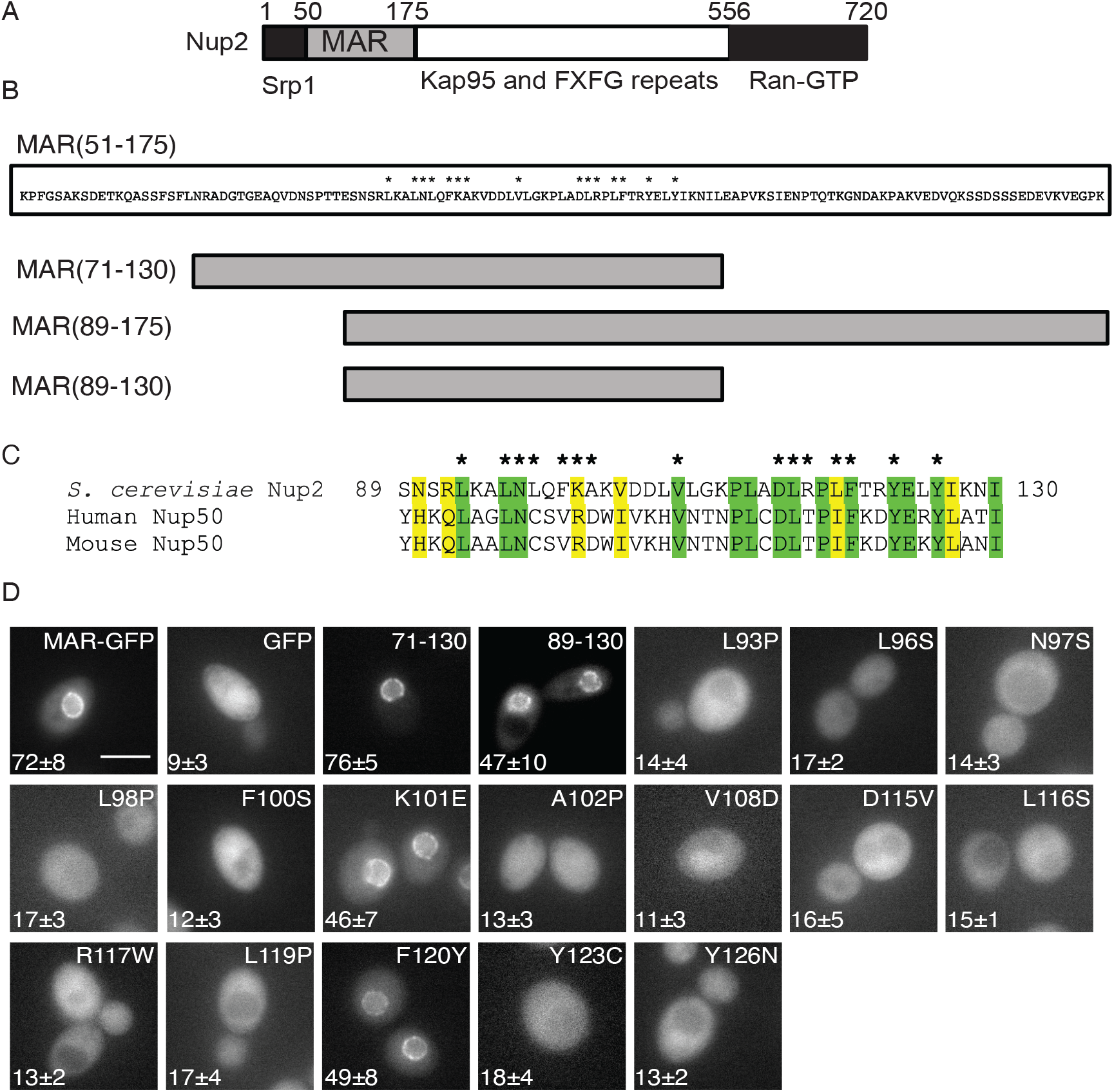
Location of point mutations in the Nup2 MAR and their effect on localization to the nuclear envelope. (A) Schematic of the Nup2 protein showing the MAR (amino acid residues 51-175) and regions of Nup2 known to bind the transport proteins Srp1, Kap95 (importin-β), and Ran. (B) Above: Amino acid sequence of the MAR, with asterisks indicating the positions of point mutations that affect MAR function. Below: Schematic depiction of deletion mutants of the MAR. (C) Alignment of the functional region of the MAR with homologous regions in human and mouse Nup50. Identical amino acids are highlighted in green, similar amino acids in yellow. Asterisks indicate positions of the MAR point mutations. (D) Representative fluorescent images of cells expressing wild-type or mutant MAR-GFP fusions. Not all images were taken on the same day. The scale bar in the wild-type MAR-GFP panel represents 5 μm. The sporulation efficiency in an *ndj1* mutant background is shown in the bottom left corner of each panel is data taken from Table 1.

A PCR-mutagenized DNA fragment encoding *MAR-GFP* was integrated into an *ndj1* strain at the *NUP2* locus. Transformants were mated to a *nup2 ndj1* strain, and the resulting diploids were screened for the ability to sporulate. Of 8700 diploids screened, 37 had reduced sporulation compared to a wild-type *MAR-GFP* strain. The mutant candidates were sequenced: 23 contained a unique point mutation, 12 were duplicates, and 2 contained more than 5 non-silent point mutations each and were excluded from further analysis. The 23 unique mutations affected 15 amino acid positions in the central portion of the MAR (Figure 1B). Diploids homozygous for the *MAR-GFP* mutations were constructed, and sporulation efficiency was measured (Table 1). Most of the mutants sporulated at 10-15% efficiency, or roughly the same as the null allele (GFP expressed from the *NUP2* promoter). The exceptions were the mutants with the amino acid substitutions K101E and F120Y, which sporulated at approximately 50% efficiency. For these two mutants, we also measured sporulation in a *mar-GFP/nup2Δ* heterozygote and found that hemizygosity decreased sporulation to around 15%.

**Table 1.** Effect of MAR mutations on sporulation

| Relevant genotype | Sporulation efficiency (n ≥ 6) |
| --- | --- |
| *MAR(51-175)-GFP*<sup>'''</sup> *ndj1Δ*<sup>'''</sup> | 72 ± 8 |
| *GFP [no MAR]*<sup>'''</sup> *ndj1Δ*<sup>'''</sup> | 9 ± 3 |
| *mar(L93P)-GFP*<sup>'''</sup> *ndj1Δ*<sup>'''</sup> | 14 ± 4 |
| *mar(L96S)-GFP*<sup>'''</sup> *ndj1Δ*<sup>'''</sup> | 16 ± 2 |
| *mar(N97D)-GFP*<sup>'''</sup> *ndj1Δ*<sup>'''</sup> | 10 ± 4 |
| *mar(N97I)-GFP*<sup>'''</sup> *ndj1Δ*<sup>'''</sup> | 12 ± 3 |
| *mar(N97S)-GFP*<sup>'''</sup> *ndj1Δ*<sup>'''</sup> | 15 ± 3 |
| *mar(N97Y)-GFP*<sup>'''</sup> *ndj1Δ*<sup>'''</sup> | 14 ± 4 |
| *mar(L98P)-GFP*<sup>'''</sup> *ndj1Δ*<sup>'''</sup> | 16 ± 3 |
| *mar(F100P)-GFP*<sup>'''</sup> *ndj1Δ*<sup>'''</sup> | 14 ± 3 |
| *mar(F100S)-GFP*<sup>'''</sup> *ndj1Δ*<sup>'''</sup> | 14 ± 4 |
| *mar(K101E)-GFP*<sup>'''</sup> *ndj1Δ*<sup>'''</sup> | 46 ± 7 |
| *mar(A102P)-GFP*<sup>'''</sup> *ndj1Δ*<sup>'''</sup> | 13 ± 3 |
| *mar(V108D)-GFP*<sup>'''</sup> *ndj1Δ*<sup>'''</sup> | 11 ± 3 |
| *mar(D115G)-GFP*<sup>'''</sup> *ndj1Δ*<sup>'''</sup> | 11 ± 3 |
| *mar(D115V)-GFP*<sup>'''</sup> *ndj1Δ*<sup>'''</sup> | 15 ± 5 |
| *mar(D115Y)-GFP*<sup>'''</sup> *ndj1Δ*<sup>'''</sup> | 14 ± 4 |
| *mar(L116S)-GFP*<sup>'''</sup> *ndj1Δ*<sup>'''</sup> | 15 ± 1 |
| *mar(R117W)-GFP*<sup>'''</sup> *ndj1Δ*<sup>'''</sup> | 13 ± 2 |
| *mar(L119P)-GFP*<sup>'''</sup> *ndj1Δ*<sup>'''</sup> | 17 ± 4 |
| *mar(F120Y)-GFP*<sup>'''</sup> *ndj1Δ*<sup>'''</sup> | 49 ± 9 |
| *mar(Y123C)-GFP*<sup>'''</sup> *ndj1Δ*<sup>'''</sup> | 14 ± 5 |
| *mar(Y123H)-GFP/" ndj1Δ/"* | 14 ± 3 |
| *mar(Y126C)-GFP/" ndj1Δ/"* | 14 ± 2 |
| *mar(Y126N)-GFP/" ndj1Δ/"* | 13 ± 2 |
| *mar(71-130)-GFP/" ndj1Δ/"* | 72 ± 8 |
| *mar(89-130)-GFP/" ndj1Δ/"* | 47 ± 10 |
| *mar(89-175)-GFP/" ndj1Δ/"* | 76 ± 4 |
| *mar(K101E)-GFP/nup2Δ ndj1Δ/"* | 15 ± 3 |
| *mar(F120Y)-GFP/nup2Δ ndj1Δ/"* | 16 ± 3 |
| *mar(89-130)-GFP/nup2Δ ndj1Δ/"* | 22 ± 3 |
| *mar(89-175)-GFP/nup2Δ ndj1Δ/"* | 72 ± 6 |

### The central region of the MAR is sufficient for its function

Because the mutations all lie in the middle of the MAR, we constructed several deletions to test whether this central fragment was sufficient to provide MAR function. A 60-amino acid stretch from amino acids 72-130 corresponding to a region of homology between Nup2 and mammalian Nup50 was sufficient to support sporulation (Table 1), indicating that amino acids 51-71 and 131-175 are dispensable for MAR function. A shorter fragment confined mostly to the region affected by the point mutations (*mar(89-130)*) was only partially functional, but because *mar(89-175)* is fully functional, it seems likely that most of the MAR function is provided by the 42-amino acid stretch from 89-130. Eleven out of the 15 affected amino acids map to conserved residues found in human and mouse Nup50 (Figure 1C).

### Most of the point mutations disrupt localization to the nuclear envelope

We next examined the mutant strains by fluorescence microscopy and found that unlike wild-type MAR-GFP, which is localized to the nuclear envelope, most of the mutants were distributed throughout the cell (Figure 1D). The K101E and F120Y fusions localized to the nuclear envelope, as did the mar(89-130)-GFP fusion. Because the mutants with proper localization sporulated with lower efficiency when hemizygous, we examined hemizygotes for these alleles and saw that the fusion localized to the nuclear envelope in all three cases (Figure S1). Thus, decreasing the copy number of these three mutants does not result in improper subcellular localization, and the lower sporulation efficiency of the hemizygotes is unlikely to be due to mis-localization.

### MAR mutants are defective in binding to Nup60

Since Nup2 is known to be recruited to the nuclear envelope by Nup60 (Hood *et al*. 2000; Solsbacher *et al*. 2000), we hypothesized that the MAR mutants with aberrant localization were defective in interacting with Nup60. To test whether the MAR mutations affect interaction with Nup60, we set up a yeast two-hybrid (Y2H) assay using the MAR as bait and Nup60 as prey. We made a Gal4 activation domain fusion to full-length Nup60 and Gal4 DNA binding domain fusions to the wild-type MAR and to a subset of the MAR mutants. While the wild-type MAR interacts with Nup60, many of the mutants do not (Figure 2). As expected, the two mutants that localize to the nuclear envelope interact with Nup60 by Y2H. However, some of the mutants that are not recruited to the nuclear envelope also interact with full length Nup60. Because the Y2H assay can detect weak or transient interactions between proteins (Mehla *et al*. 2017), we suspected that a defect in the interaction of these mutants with Nup60 could be masked if the MAR is able to interact with more than one part of Nup60.

**Figure 2.**
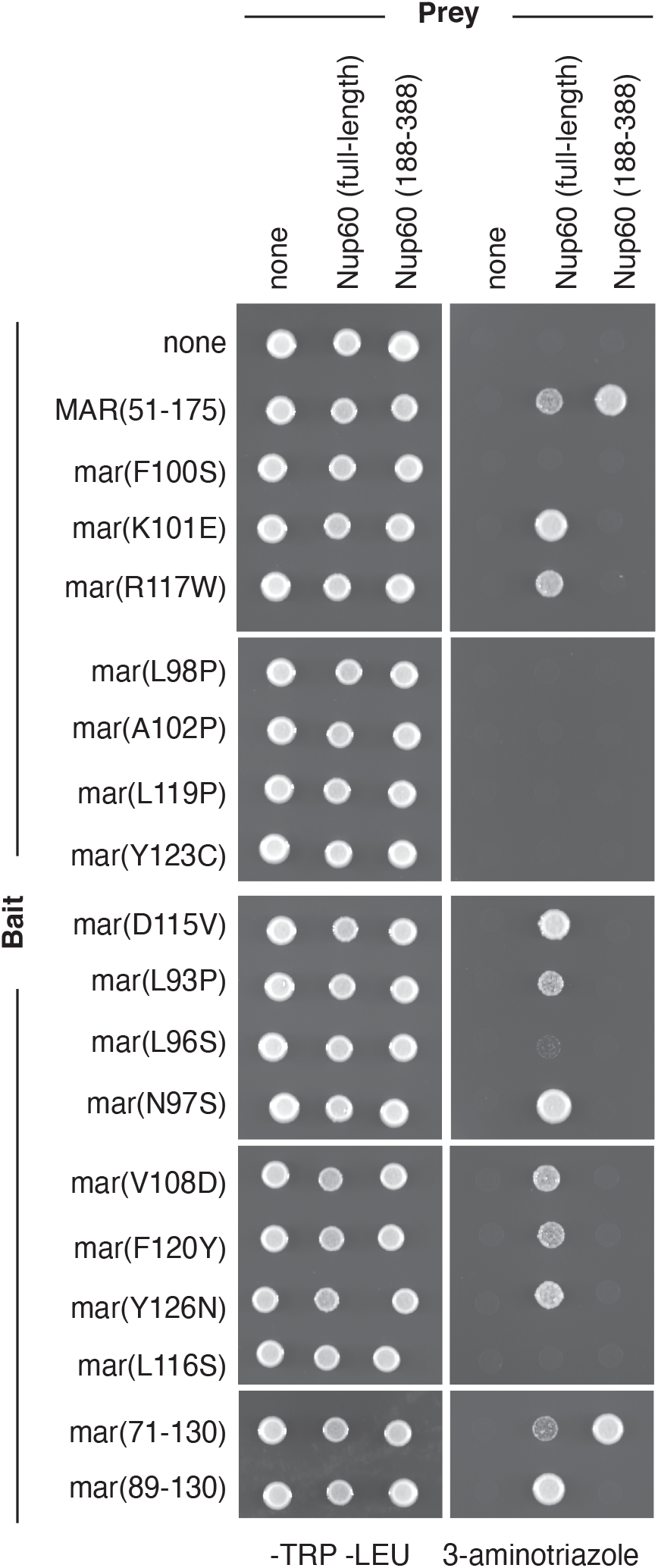
Interaction of MAR mutants with Nup60 in the Y2H assay. Panels show yeast two-hybrid assays of wild-type and mutant MARs interacting with either full-length Nup60 or Nup60(188-388). A pGAL1-HIS3 strain transformed with bait and prey plasmids was spotted onto -TRP-LEU (left panels) and -TRP-LEU-HIS + 5 mM 3-aminotriazole plates (right panels). The MAR alleles used as bait are indicated on the left-hand side; the fragments of Nup60 used as prey are indicated on the top. Bait controls were included on all the plates but are only shown for the top panels.

Pull down assays with purified fragments of Nup2 and Nup60 have shown that Nup2 interacts with both an N-terminal fragment of Nup60 from amino acids 1-188 and an internal fragment of Nup60 from amino acids 189-388 (Denning *et al*. 2001). Although most of the Nup60 fragments we tested failed to interact with the wild-type MAR in the Y2H assay, we were able to detect an interaction between Nup60(188-388) and the MAR. Furthermore, all the MAR mutations disrupt this interaction, including the mutations that do not affect localization to the nuclear envelope. These results indicate that all the mutations affect interaction with Nup60 to some degree, although the defect is not always severe enough to prevent the MAR from binding to Nup60 at the nuclear envelope.

### Binding of the MAR to meiotic chromosomes requires Nup60

We next tested whether the mutant MAR-GFP fusions bind to meiotic chromosomes by staining chromosome spreads with an antibody against GFP. While wild-type MAR-GFP forms foci on meiotic chromosomes, the mutants were absent from the spreads (Figure 3A). Quantitation of the fluorescent signal showed that binding of the mutants was comparable to that of GFP (Figure 3B). Therefore, the mutants appear to be defective in interacting with both Nup60 and meiotic chromosomes, suggesting that the MAR might require Nup60 to bind meiotic chromosomes.

**Figure 3.**
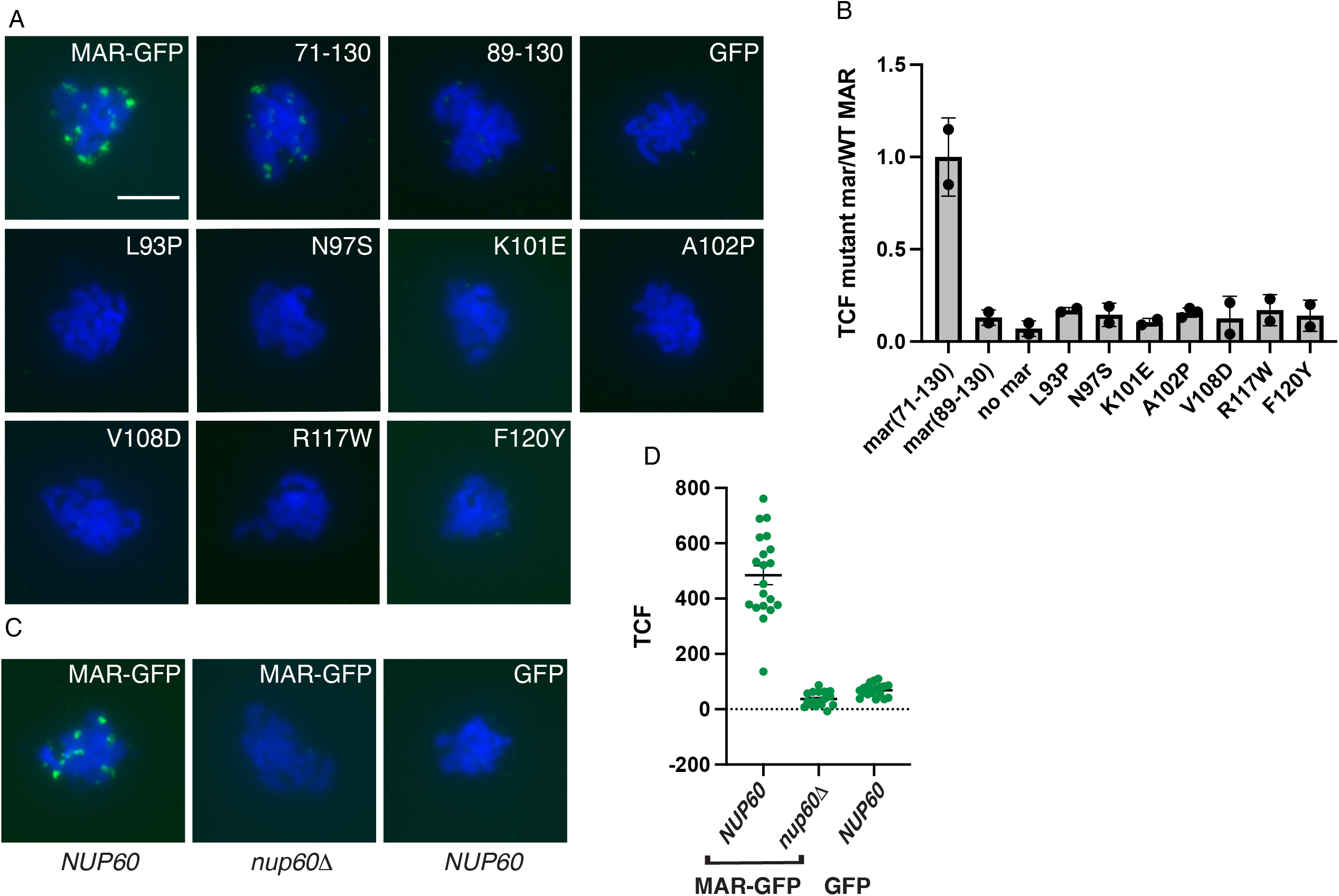
Effect of MAR mutations on binding to meiotic chromosomes. (A) Representative meiotic chromosome spreads showing the localization of wild-type and mutant MAR-GFP fusions. Cells were harvested 10 hours after transfer to sporulation medium then spread onto glass slides. The MAR-GFP fusions were detected using a polyclonal antibody to GFP and an Alexa Fluor 488-conjugated secondary antibody. The scale bar in the wild-type panel represents 2 μm. (B) Mean relative fluorescence intensities of mutant MAR-GFP fusions normalized to wild-type MAR controls done on the same day. Each data point is the average fluorescence intensity from at least 10 nuclei. Measurements from day-matched individual cells are shown in Figure S3. P<0.0001 for all mutant/WT comparisons using two-way ANOVA tests except for mar(71-130) in which P=0.95. (C) Representative meiotic chromosome spreads showing the localization of wild-type MAR-GFP in *NUP60* and *nup60*Δ strains and of GFP in a *NUP60* strain. The scale bar in the left panel represents 5 μm. (D) Quantitation of total corrected fluorescence of MAR-GFP from spreads represented in (C). P<0.0001 using a two-tailed t-test.

To test whether the MAR can bind to meiotic chromosomes in the absence of Nup60, we prepared chromosome spreads from *NUP60* and *nup60Δ* strains expressing wild-type MAR-GFP and stained the spreads with antibodies against GFP (Figure 3C). Binding of MAR-GFP in the *nup60Δ* strain was reduced to the background levels exhibited by GFP alone (Figure 3D). These results demonstrate that the MAR requires Nup60 to bind to meiotic chromosomes.

### Loss of Nup60 blocks sporulation in strains with telomere bouquet defects

The observation that Nup60 was required for binding of the MAR to meiotic chromosomes led us to speculate that Nup60 might be playing a larger role in promoting sporulation than we initially suspected. To explore this possibility, we began by asking whether *nup60Δ*, like *nup2Δ*, blocks sporulation in strains lacking *NDJ1* or *CSM4*. Both *nup60Δ* and *nup60Δndj1Δ* strains grow poorly compared to wild-type strains; however, only the double mutant has a significant sporulation defect (Figure 4A and B, row 1), indicating that the growth defect alone does not result in poor sporulation and that *nup60Δ* is synthetic with *ndj1Δ*. Likewise, the *nup60Δcsm4Δ* strain exhibits a synthetic sporulation defect, albeit to a lesser degree than the *nup2Δcsm4Δ* double mutant (Table 2). We predicted that *nup60Δ* and *nup2Δ* should not have a synthetic phenotype if the two proteins are acting in concert to promote sporulation. However, because the *nup60Δnup2Δ* double mutant either grows extremely poorly or is inviable depending on the strain background (Dilworth *et al*. 2001), epistasis analysis required an allele of *NUP60* that affected sporulation but not growth.

**Figure 4.**
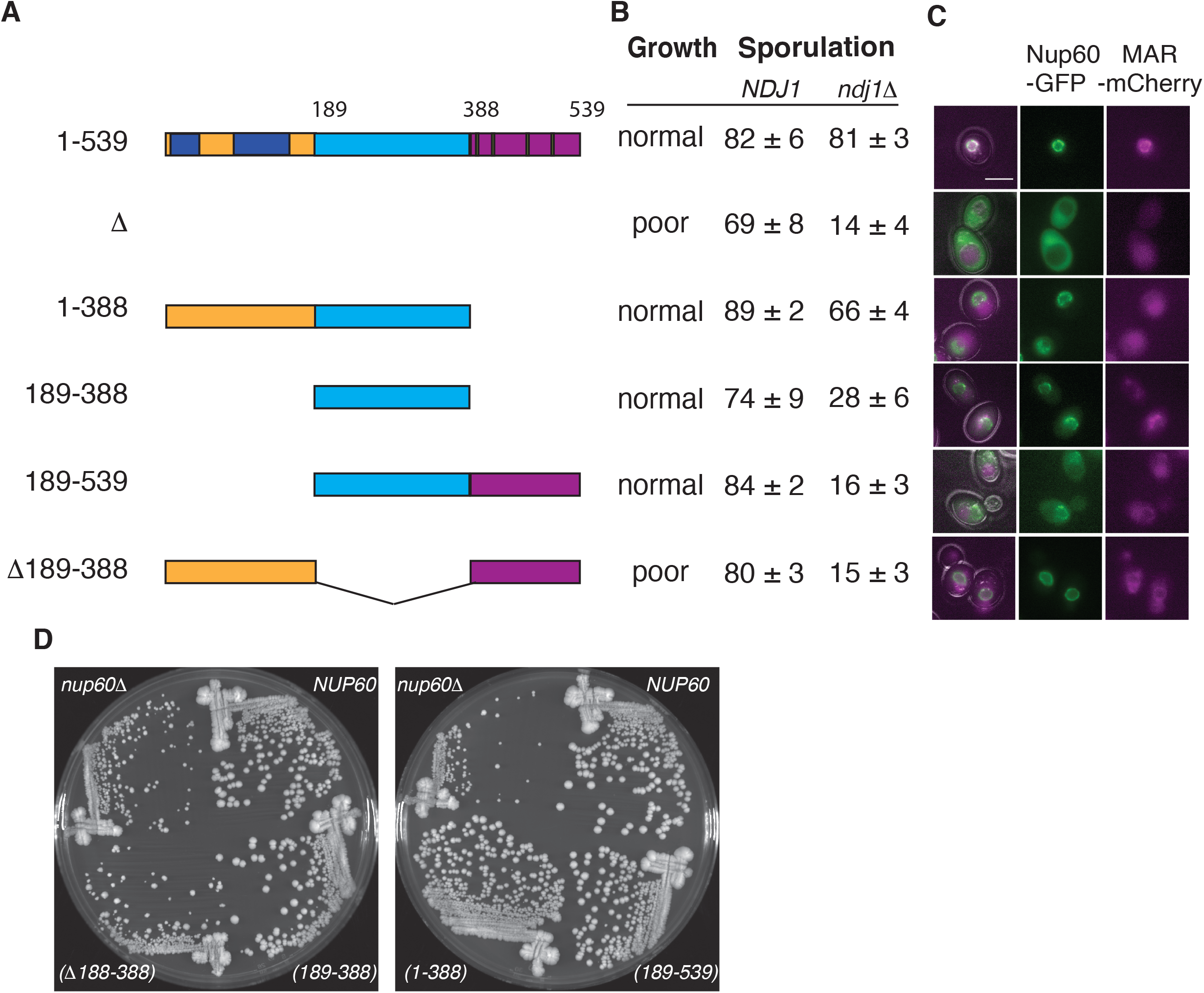
Effect of Nup60 truncations on growth, sporulation, and recruitment of the MAR to the nuclear envelope. (A) Schematic representation of full length Nup60 and truncation mutants. Full-length Nup60 contains an N-terminal region from amino acids 1-188 (yellow), a middle region from aa 189-388 (light blue), and a C-terminal region from aa 389-539 (purple). Two helices that help anchor Nup60 to the nuclear envelope are in dark blue; four FxF repeats are in green. (B) Table showing growth and sporulation of the *NUP60* mutants illustrated in (A). All strains in the growth and *NDJ1* sporulation column are isogenic to the wild-type (1-539) strain: *NUP60(1-539)* (SBY1903), *nup60Δ* (SBY5217), *nup60(1-388)* (SBY6423), *nup60(189-388)* (SBY6420), *nup60(189-539)* (SBY6426), *nup60(Δ189-388)* (SBY6429). All strains in the *ndj1* column are isogenic to the *NUP60(1-539)* strain Δ (SBY1904): *nup60* (SBY5222), *nup60(1-388)* (SBY6296), *nup60(189-388) (SBY6293), 189-539* (SBY6299), *nup60(*Δ*189-388)* (SBY6302). (C) Representative images of cells expressing wild-type MAR-mCherry and GFP fusions to the Nup60 truncations illustrated in (A) Intensity of the *NUP60(1-539)* MAR-mCherry panel was adjusted to highlight its localization to the NE. The scale bar represents 5 μm. All strains are isogenic to the *NUP60(1-539)-GFP* strain: *(1-539)* (SBY6254),*Δ*(SBY6338), *(1-388)* (SBY6344), *(189-388)* (SBY6341), *(189-539)* (SBY6347), *(Δ189-388)* (SBY6350). (D) Colony growth of *NUP60* truncation mutants on YPD plates. Strains names are listed above in (B).

**Table 2.** *nup60* is synthetic with *csm4* but not with *nup2*

| Relevant genotype | Sporulation efficiency (n ≥ 6) |
| --- | --- |
| Wild-type | 82 ± 6 |
| *nup60Δ*<sup>'''</sup> | 69 ± 8 |
| *nup60(189-388)*<sup>'''</sup> | 74 ± 9 |
| *csm4Δ*<sup>'''</sup> | 80 ± 5 |
| *nup60Δ*<sup>'''</sup> *csm4Δ*<sup>'''</sup> | 15 ± 3 |
| *nup60(189-388)*<sup>'''</sup> *csm4Δ*<sup>'''</sup> | 13 ± 8 |
| *nup2Δ*<sup>'''</sup> *csm4Δ*<sup>'''</sup> | 6 ± 2 |
| *nup2Δ*<sup>'''</sup> | 82 ± 5 |
| *nup2Δ*<sup>'''</sup> *nup60(189-388)*<sup>'''</sup> | 74 ± 7 |
| *nup2Δ*<sup>'''</sup> *ndj1Δ*<sup>'''</sup> | 15 ± 4 |
| *nup2Δ*<sup>'''</sup> *ndj1Δ*<sup>'''</sup> *nup60(189-388)*<sup>'''</sup> | 18 ± 5 |

### A Nup60 N-terminal fragment is sufficient for sporulation function

We constructed a set of *NUP60* truncations with the aim of finding an allele defective for sporulation alone. The N-terminus of Nup60 contains two helical regions that contact the nuclear envelope and the NPC core, while the C-terminus contains FxF repeats that bind nuclear transport receptors (Denning *et al*. 2001; Mészáros *et al*. 2015). We hypothesized that deleting the central portion of Nup60 would create the desired allele by removing a Nup2 interaction domain while leaving regions involved in pore structure and nuclear transport intact. Instead, we found that the strain missing the central domain of Nup60 (Δ189-388) grows as poorly as the *nup60Δ* mutant, while another expressing only the central portion of Nup60 (189-388) grows as well as wild type (Figure 4B and D). In addition, we tested growth on hydroxyurea (HU) plates, since *nup60Δ* strains are known to be hypersensitive to DNA-damaging agents (Bermejo *et al*. 2011). The Nup60(189-388) fragment was necessary and sufficient for wild-type growth in the presence of HU (Figure S2).

We next tested how the *NUP60* deletions affect sporulation and found that only full-length Nup60 and the N-terminal fragment from amino acids 1-388 complement the sporulation defect (Figure 4B). Although the middle fragment from 189-388 fully rescues the growth defect, it complements the sporulation defect only slightly. Collectively, these results demonstrate that the middle region of Nup60 is necessary but not sufficient for sporulation, while the C-terminal region is completely dispensable.

### *NUP60* and *NUP2* are in the same genetic pathway

The ability of the *nup60(189-388)* allele to complement growth but not sporulation allowed us to test for a synthetic interaction between *nup2* and *nup60*. We found that *nup60(189-388)* on its own, like *nup2*, has little effect on sporulation and that the *nup2 nup60* double mutant also sporulates with wild-type efficiency (Table 2). In addition, the *nup60(189-388) nup2 ndj1* triple mutant sporulates with approximately the same efficiency as either the *nup60(189-388) ndj1* or *nup2 ndj1* double mutant. Thus, *nup60(189-388)* and *nup2* do not have a synthetic sporulation phenotype. We also measured the kinetics of nuclear division in a meiotic time course of wild-type, *nup2Δ, nup60(189-388)*, and *nup2Δ nup60(189-388)* strains. Again, *nup60(189-388)* phenocopies *nup2Δ*, causing an approximately one-hour delay in the first meiotic division, while the double mutant exhibits a delay similar to that seen in the single mutants (Figure 5A). In addition, we compared the kinetics of nuclear division in wild-type, *ndj1Δ, nup60(189-388)* and *ndj1Δ nup60(189-388)* strains. The *ndj1Δnup60(189-388)* double mutant takes significantly longer to undergo the first meiotic division than either single mutant (Figure 5B).

**Figure 5.**
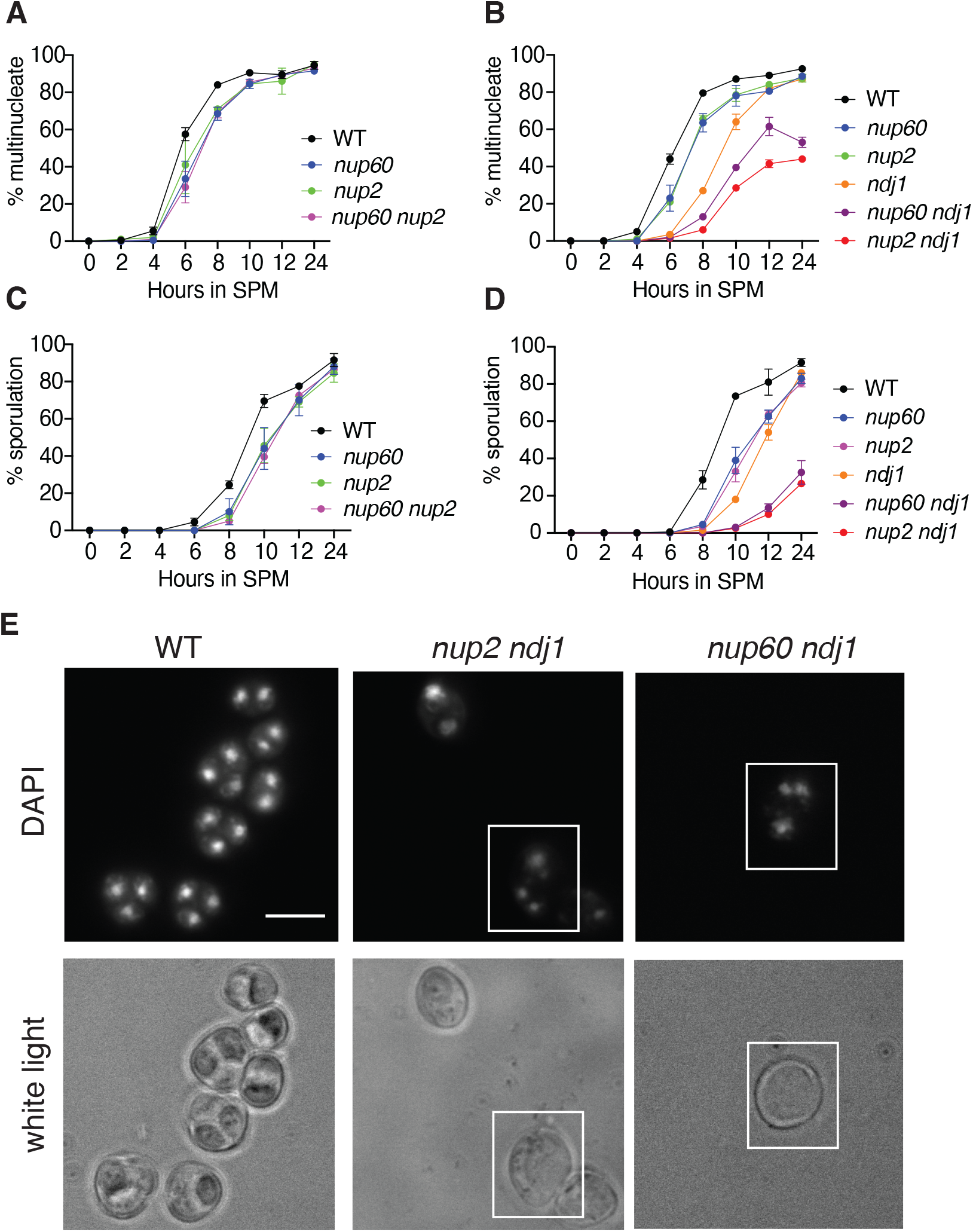
Effect of *nup60(189-388)* on the kinetics of nuclear division. (A) Epistasis analysis of *nup60(189-388)* and *nup2*. Synchronized cells cultured in liquid sporulation medium were removed for analysis at the indicated times after initial transfer. At least 200 cells were analyzed for the presence of either one or more than one DAPI-staining body. All strains are isogenic to the wild-type strain: wild-type (black; SBY1903) *nup60* (blue; SBY6293), *nup2* (green; SBY3945), and *nup60 nup2* (magenta; SBY6432). A representative time course is shown for four strains tested in duplicate on the same day. (B) Epistasis analysis of *nup60(189-388)* and *ndj1*. A time course experiment was performed as described in (A) using isogenic strains: WT (black; SBY1903), *nup60* (blue; SBY6293), *nup2* (green; SBY3945), *ndj1* (orange; SBY1904), *nup60 ndj1* (purple; SB6296), and *nup2 ndj1* (red; SBY3983). A representative time course is shown for six strains tested in duplicate on the same day. (C) Sporulation counts of samples from the time course depicted in (A). For each sample, at least 200 cells were analyzed under white light for the presence of two or more spores. (D) Sporulation counts of samples from the time course depicted in (B). (E) Images of DAPI-stained samples of wild-type, *nup2 ndj1*, and *nup60 ndj1* strains after 24 hours in sporulation media. The top panels show DAPI fluorescence, and the bottom panels show the same cells under white light. The white boxes highlight examples of cells where division of the nuclear masses has taken place without spore formation.

These results demonstrate that *NUP60* acts in the same pathway as *NUP2* but in a different pathway from *NDJ1*. Strains that showed a delay in the first meiotic division also exhibited a delay in spore formation (Figures 5C, D). Oddly, in the *nup2 ndj1* and *nup60 ndj1* strains, the percentage of multinucleate cells is significantly higher than the percentage of sporulated cells at the 24-hour time point. When we examined the DAPI stained cells under white light, we noticed that many of the multinucleate cells in these double mutants had not formed visible spores (Figure 5E). In contrast, when we examined the wild-type strain, all multinucleate entities examined had sporulated.

### Full-length Nup60-GFP is required to recruit the MAR to the NE

To determine which part or parts of Nup60 can recruit the MAR to the nuclear envelope, we tagged the Nup60 fragments with GFP. We first ascertained that the GFP-tagged alleles behave like their untagged counterparts in terms of growth and sporulation (Figure S4). We next examined strains expressing each of the tagged Nup60 fragments by fluorescence microscopy and found that all the Nup60-GFP fusions localized to the nuclear envelope to some degree (Figure 4C, middle column). However, fusions lacking the N-terminal 188 amino acids were also aberrantly present in the nucleoplasm and cytoplasm. These results are consistent with published observations that helical domains in the N terminus of Nup60 are important for contacting the nuclear membrane and the core of the NPC (Mészáros *et al*. 2015).

When we examined the localization of MAR-mCherry that was co-expressed in each of the Nup60-GFP strains, we found that MAR-mCherry is properly localized at the nuclear envelope only in the strain expressing full-length Nup60-GFP (Figure 4C, right column). When the N-terminus, middle, or C-terminus of Nup60 is deleted, MAR-mCherry appears largely throughout the cell. Curiously, Nup60(1-388)-GFP supports sporulation despite being unable to recruit the MAR to the nuclear envelope.

### Full-length Nup60-GFP is required to recruit the MAR to meiotic chromosomes

We next examined binding of the Nup60 fragments to meiotic chromosomes spreads. We found that the only Nup60 fragment that supports sporulation (Nup60(1-388)) binds to meiotic chromosomes while the Nup60 fragments that do not support sporulation do not bind to meiotic chromosomes (Figure 6A and B, middle column). These results show that the N-terminal fragment from amino acids 1-388 is sufficient for meiotic chromosome binding and demonstrate that there is a correlation between Nup60’s chromosome-binding activity and its ability to promote sporulation.

**Figure 6.**
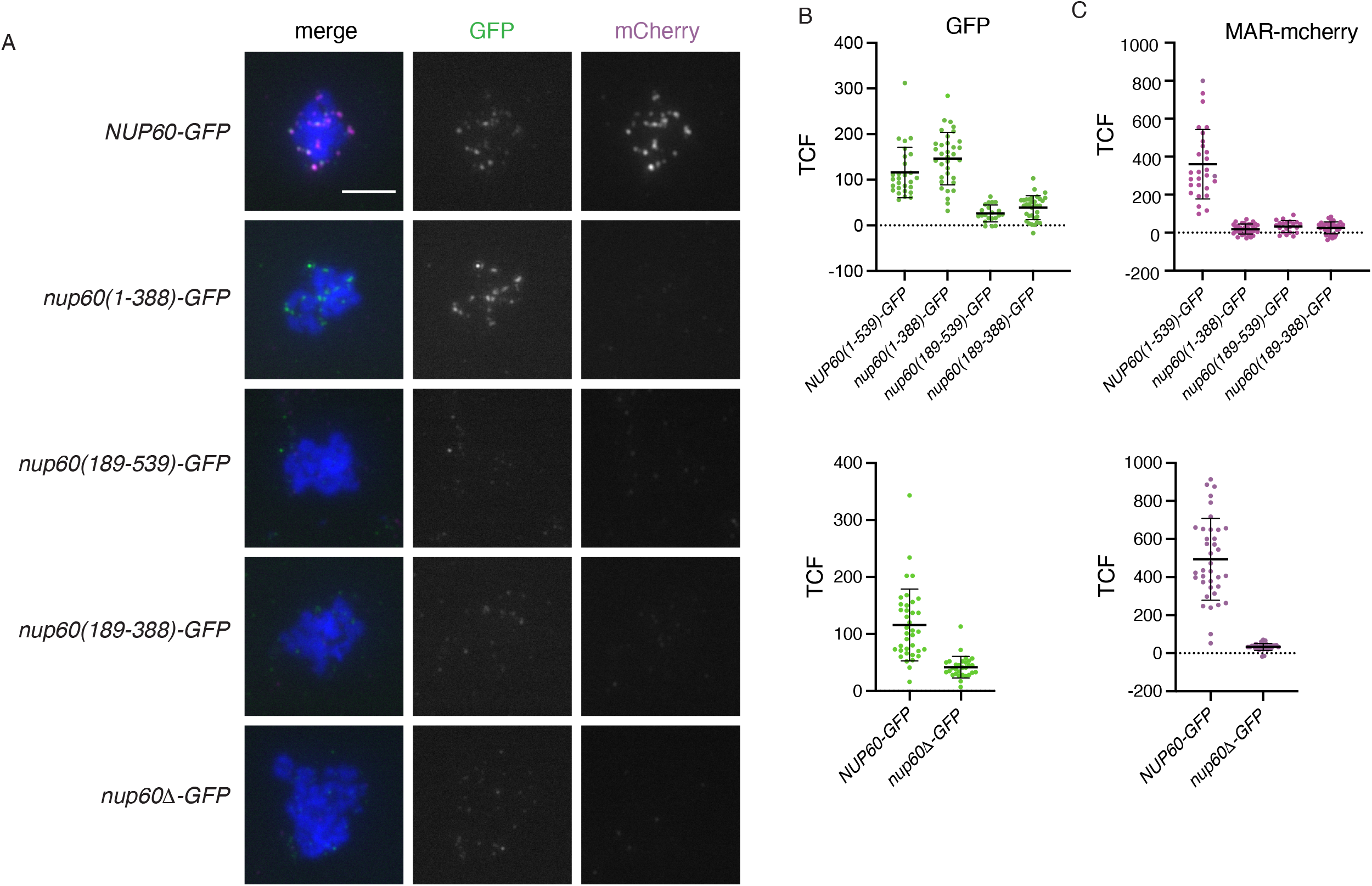
Effect of Nup60 truncations on recruitment of the MAR to meiotic chromosomes. (A) Representative meiotic chromosome spreads from strains expressing MAR-mCherry and full-length or truncated Nup60-GFP fusions. Cells were harvested 10 hours after transfer to sporulation medium then spread onto glass slides. The Nup60-GFP fusions were detected using a polyclonal antibody to GFP and an Alexa Fluor 488-conjugated secondary antibody; MAR-mCherry was detected using a polyclonal antibody to mCherry and an Alexa Fluor 594-conjugated secondary antibody. The scale bar in the upper left panel represents 5 μm. All strains are isogenic to the *NUP60-GFP* strain: *NUP60-GFP* (SBY6254), *nup60(1-388)-GFP* (SBY6344), *nup60(189-388)-GFP* (SBY6341), *nup60(189-539)-GFP* (SBY6347); *nup60Δ-GFP* (SBY6338) was spread and analyzed on a different day (WT not shown to save space). (B) Quantitation of total corrected fluorescence of Nup60-GFP from spreads represented in (A). P=0.0125, comparing *NUP60(1-539)* and *nup60(1-388)*; P<0.0001 for all other comparisons to (1-539) using one-way ANOVA with Dunnett’s multiple comparisons tests. (C) Quantitation of total corrected fluorescence of MAR-mCherry from spreads represented in (A). P<0.0001 for all comparisons to (1-539) using the same method as in (B).

In spreads of strains expressing full-length Nup60-GFP, both Nup60-GFP and MAR-mCherry are present on the chromosomes and appear to be co-localized with one another. In contrast, in spreads with Nup60 fragments that do not bind to the chromosomes, MAR-mCherry is absent (Figure 6A and C, right column). Finally, although Nup60(1-388)-GFP is present on meiotic chromosomes, it does not appear to recruit MAR-mCherry. These results indicate that the C-terminus of Nup60 is dispensable for the binding of Nup60 to meiotic chromosomes but required for binding to the MAR. Since Nup60(1-388) is sufficient for sporulation but does not appear to interact with the MAR at the nuclear envelope or on meiotic chromosomes, it seemed likely that Nup60(1-388) does not require Nup2 to function.

### Truncated Nup60 promotes sporulation independently of Nup2

To test whether deleting the C-terminus of Nup60 alleviates the requirement for Nup2 in sporulation, we compared *nup2Δndj1Δ* diploids expressing full-length Nup60, Nup60(1-388), or Nup60(1-388)-GFP. While the *NUP60* strain sporulates poorly, the *nup60(1-388)* and *nup60(1-388)-GFP* diploids both sporulate nearly as well as a strain that is completely wild type (Table 3). Oddly, we found that the full length Nup60-GFP fusion also restored sporulation. The nearly wild-type level of sporulation observed with the *NUP60-GFP nup2 ndj1* strain is in sharp contrast to the poor sporulation of the *nup2 ndj1* strain in which Nup60 is untagged. To test whether the GFP tag itself was conferring a new function onto full-length Nup60, we also measured sporulation in *nup2 ndj1* strains where Nup60 was tagged with mCherry or the myc antigen. Since *NUP60-mCherry* and *NUP60-13myc* also suppress the *ndj1 nup2* sporulation defect, it seems more likely that the tags are somehow interfering with the C-terminus of Nup60, causing the full-length fusions to behave like the truncated alleles. In addition, we found that suppression of the *nup2 ndj1* sporulation defect is dominant, indicating that the presence of untagged, full-length Nup60 does not prevent the tagged or truncated versions from functioning in the absence of Nup2.

**Table 3.**
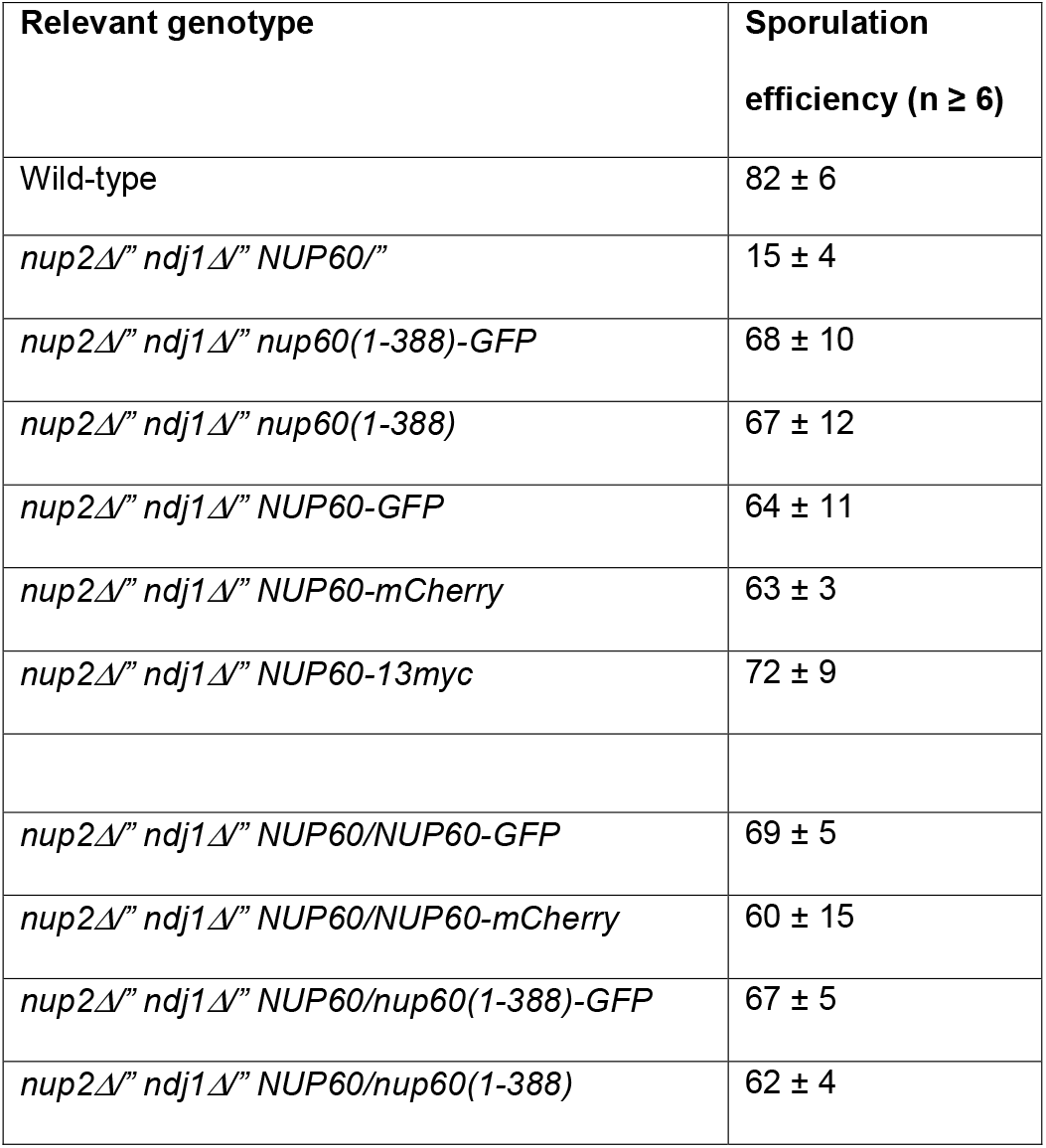
Disruption of the Nup60 C-terminus suppresses the *nup2 ndj1* sporulation defect

### The Nup60-GFP fusion binds to meiotic chromosomes in the absence of Nup2

Although Nup60-GFP is able to recruit MAR-mCherry to both the nuclear envelope and meiotic chromosome spreads, the efficient sporulation of the *NUP60-GFP ndj1 nup2* strain suggested that the Nup60-GFP fusion does not actually require the presence of the MAR to bind to meiotic chromosomes. We prepared meiotic spreads from *NUP60-GFP* and *NUP60-GFP nup2* strains and found that Nup60-GFP was bound to the chromosomes in both cases (Figure 7). Therefore, Nup60-GFP is capable of binding to meiotic chromosomes without Nup2.

**Figure 7.**
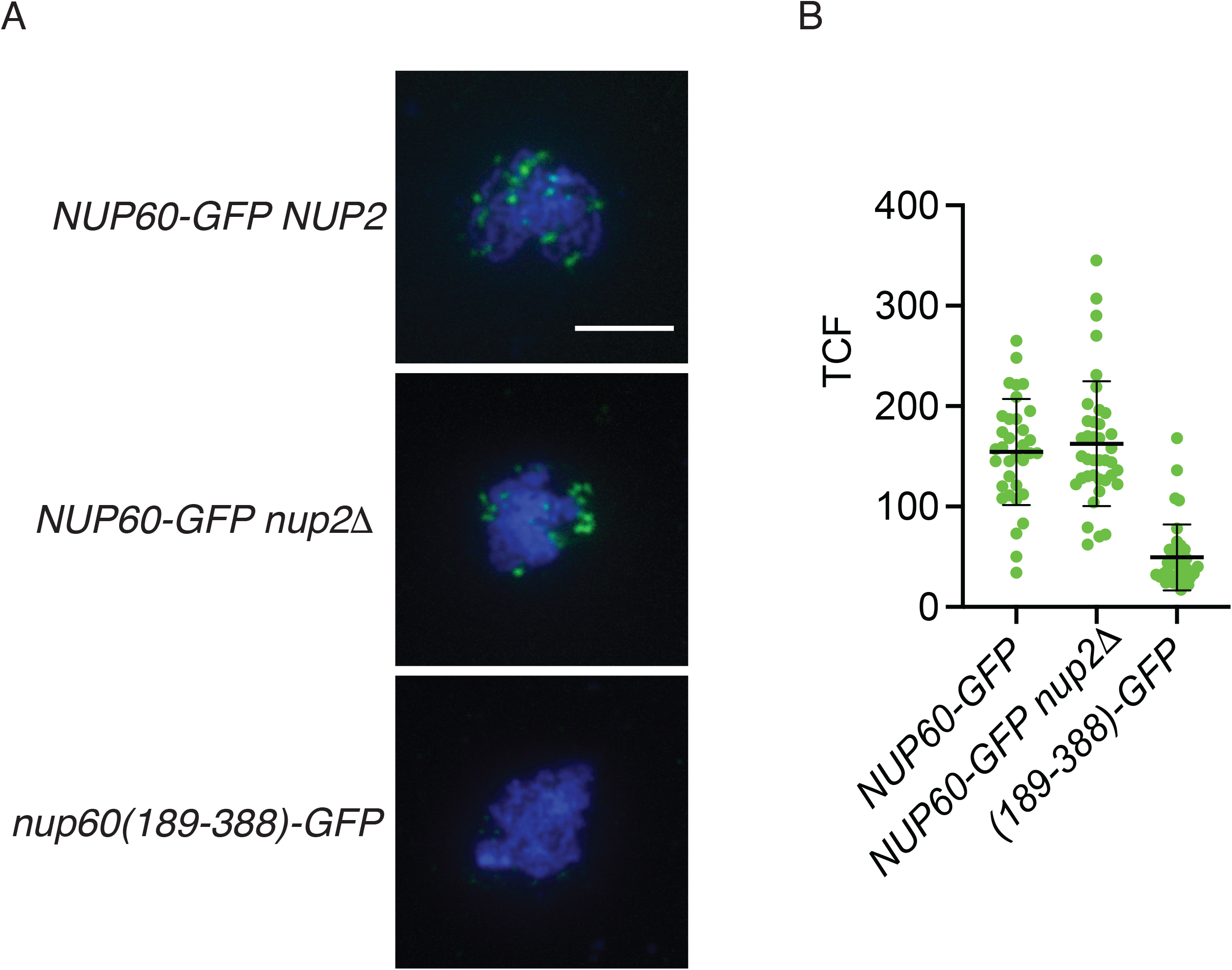
Binding of Nup60-GFP to meiotic chromosomes in the absence of Nup2. (A) Representative meiotic chromosome spreads from wild-type (SBY6305) or *nup2* (SBY6487) strains expressing Nup60-GFP. Nup60(189-388)-GFP (SBY6311) was used as the negative control because chromosome binding is comparable to that of GFP, but the cells have wild-type growth. The Nup60-GFP fusions were detected using a polyclonal antibody to GFP and an Alexa Fluor 488-conjugated secondary antibody. The scale bar in the upper left panel represents 5 μm. (B) Quantitation of total corrected fluorescence of Nup60-GFP from spreads represented in (A). P=0.5477, comparing Nup60-GFP in wild-type versus *nup2* using a two-tailed t-test.

## Discussion

Previously, we showed that a mutation in *NUP2* produces a synthetic sporulation phenotype when combined with a mutation in either *NDJ1* or *CSM4*, two genes required for the association of meiotic telomeres with the LINC complex. These results suggested that the NPC and the LINC complex contribute to sporulation through functionally redundant but distinct pathways, perhaps by organizing meiotic chromosomes in the nucleus. The work presented here builds on those earlier findings to show that mutations in *NUP60* also give a synthetic sporulation defect with *ndj1Δ* and *csm4Δ* mutations. Moreover, while binding of the Nup2 MAR to both the nuclear envelope and meiotic chromosomes is completely dependent on Nup60, the N-terminus of Nup60 is capable of binding chromosomes and promoting sporulation in the absence of Nup2. This is significant because Nup2 is thought to be only transiently associated with the NPC, while Nup60’s interaction with the core NPC is more stable. If Nup60 rather than Nup2 is responsible for promoting sporulation, that would suggest that the association of chromatin with the nuclear envelope is important for sporulation.

Normally, both Nup60 and the Nup2 MAR are required for sporulation in *ndj1* mutants: it is only when the C-terminus of Nup60 is deleted that Nup2 is no longer needed. These results indicate that the sporulation function of Nup60 can be attributed to the N-terminal domain, that the C-terminus somehow represses this function, and that Nup2 counteracts that inhibition (Figure 8). One simple explanation for how autoinhibition of Nup60 might occur is that the protein interacts with itself and that binding of the MAR disrupts this interaction. AlphaFold modeling of Nup60’s structure (Jumper *et al*. 2021) predicts that the Nup60 C-terminus could potentially mask a large part of the N-terminal domain (Figure S5B). Although the confidence score is very poor for most of the Nup60(389-539) fragment, and FxF repeat regions are thought to be largely unstructured (Denning *et al*. 2003), the helix at the extreme C-terminus is predicted with high confidence and may be responsible for mediating an interaction between Nup60’s N- and C-terminal domains.

**Figure 8.**
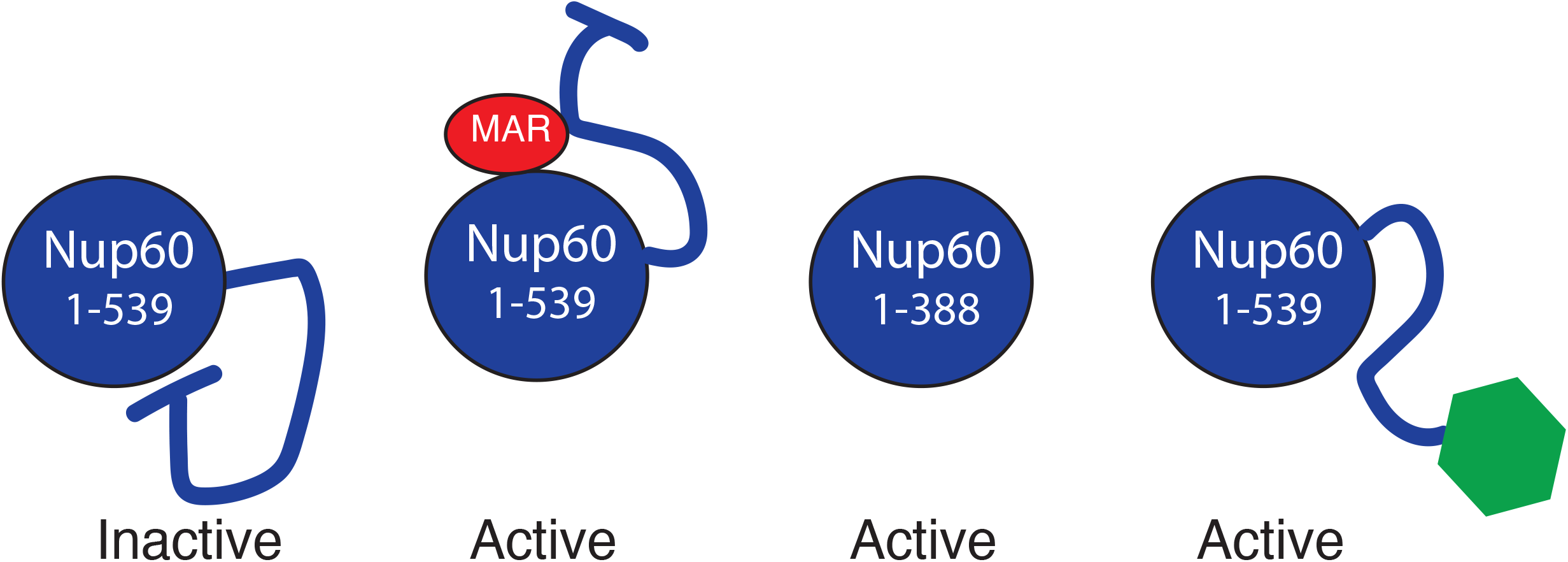
Cartoon representation of Nup60 auto-inhibition. Left: In the absence of Nup2, Nup60 is inactive due to inhibition by its own C-terminus. Middle-left: Binding of the Nup2 MAR relieves this inhibition. Middle-right: When the C-terminus is deleted, Nup60 is active, even in the absence of Nup2. Right: Fusion of GFP to the end of Nup60 prevents the C-terminus from inhibiting Nup60 function. The cartoon is meant to depict functional interactions between the MAR and regions of Nup60 and are not necessarily representative of actual physical interactions.

Another possibility is that an inhibitory protein binds to the Nup60 C-terminus either directly or as cargo of the transport machinery. Fusion of a fluorescent protein (GFP, mCherry) or an epitope tag (13myc) to Nup60’s C-terminus mimics the effect of a C-terminal deletion, as any of these modifications disrupts Nup60’s autoinhibition. A parallel phenomenon has been observed with Nup1, where adding a large protein tag or deleting the C-terminal 36 amino acids interferes with Nup1’s ability to recruit damaged telomeres to the NPC but does not affect any of Nup1’s other known functions (Aguilera *et al*. 2020). Like Nup60, Nup1 is considered to be a functional homolog of mammalian Nup153, partly due to the presence of a conserved importin-α docking site at the extreme C-termini of Nup1 and Nup153, which in Nup1 is known to bind importin-β (Kap95) (Pyhtila and Rexach 2003; Sistla *et al*. 2007). While the predicted helix at the end of Nup60 is not homologous to the Nup1/Nup153 docking site, importin-α/ β heterodimers bind to Nup60 via its C-terminus, and this binding can prevent Nup60 from interacting with Nup2 (Denning *et al*. 2001).

If the main role of Nup2 in promoting sporulation is to counteract the inhibitory effect of the Nup60 C-terminus, then it is perhaps not surprising that only a short segment of Nup2 is required for function and that the mutations identified by our screen all seem to affect the ability of the MAR to bind to Nup60. In principle, inhibition could be achieved simply by masking a small surface on Nup60. While a crystal structure of full-length Nup2 is not available, AlphaFold predicts that Nup2 contains two helices that span residues 87-132 (Jumper *et al*. 2021) and coincide almost exactly with the functional region of the MAR (residues 89-130) (Figure S5C). When we map the MAR mutations identified by our screen onto these modeled helices, we find that many of the mutations lie on the side of the helices facing a groove in the Nup2 protein (Figure S5D), potentially defining a surface of the MAR that interacts with Nup60. Most of the remaining mutations map to the interface between the two helices and possibly result in partial unfolding of the MAR.

We identified three MAR mutants (K101E, F120Y, and mar(89-130)) that localize properly to the nuclear envelope but do not bind to meiotic chromosomes or promote sporulation. Despite their proper subcellular localization, there is evidence that these mutations also alter the MAR’s interaction with Nup60. First, the three mutants do not interact with the central fragment of Nup60 by Y2H. Second, the sporulation phenotype is severe in hemizygous (*mar/Δ*) strains but mild in *mar/mar* homozygotes, suggesting that the defect can be overcome by increasing the amount of mutant MAR protein. Finally, K101E and F120Y lie on the same face of the MAR helices as most of the mutations that disrupt localization to the nuclear envelope. One possibility is that K101E, F120Y and mar(89-130) bind to Nup60 but not in a way that counteracts the inhibitory effect of the C-terminus; as a result, the mutant MAR/Nup60 complex is unable to interact with meiotic chromosomes.

Because C-terminally tagged Nup60 can bind to meiotic chromosomes without Nup2, it is tempting to speculate that the function of the C-terminus is to inhibit chromosome binding and that Nup2 overcomes that inhibition. However, it is important to point out that we do not yet know whether untagged Nup60 binds to meiotic chromosomes in the absence of Nup2. A Nup60 antibody is not commercially available, and we have been unable to construct a tagged version of Nup60 that behaves like the untagged protein despite numerous attempts (Table 3, Figure S6). We believe that the chromosome binding exhibited by Nup60-GFP fusion reflects the intrinsic chromosome binding activity of untagged Nup60, as Nup60-mCherry and Nup60-3HA also bind to meiotic chromosomes (Figure S7).

We note that the results presented here differ from our previously published work in two respects. First, the sporulation efficiencies are slightly lower for wild type (∼90% vs. 100%) and higher for *ndj1 nup2* (10% vs. 1%). Second, whereas we previously saw almost no meiotic division (i.e., separation of chromosome masses at meiosis I) in the *ndj1 nup2* double mutant, we now see that a significant proportion of cells (∼ 40%) eventually execute both meiotic divisions. We see these post-meiotic cells both in newly constructed strains and in strains that were used in Chu et al., 2017. We do not yet have a full explanation for the disparity. However, the new observations do not substantially alter the basic conclusion that loss of Nup2 causes a meiotic delay that is made more severe by the absence of Ndj1. The formation of multinucleate cells in *ndj1 nup2* strains is also consistent with the previous observation that the double mutant undergoes two rounds of spindle-pole body duplication, suggesting that meiotic progression is taking place (Chu et al., 2017).

Although a significant fraction of *nup2 ndj1* and *nup60 ndj1* cells execute both meiotic divisions, many of these cells fail to form visible tetrads. This implies that defects in the NPC basket affect both meiosis and the sporulation process itself. One possibility is that an extended delay in meiosis I somehow interferes with proper sporulation, perhaps by interfering with NPC assembly following meiotic rejuvenation. During the second division of budding yeast meiosis, rDNA circles, nucleolar proteins, and core nucleoporins of the mother cell are sequestered away from the four gamete nuclei into a fifth vesicle that is ultimately excluded from the spores (Fuchs and Loidl 2004; King *et al*. 2019). This process eliminates age-damaged nuclear components but leaves the spore nuclear membranes bereft of intact NPCs. The basket nucleoporins are the only NPC subunits that segregate with the chromatin. While neither Nup2 nor Nup60 has been directly implicated in NPC assembly, their metazoan homologs Nup50 and Nup153 have been shown to seed interphase NPC assembly in vitro (Schwartz *et al*. 2015; Vollmer *et al*. 2015). Because the yeast nuclear membrane remains intact throughout mitosis and meiosis, post-meiotic NPC assembly most likely resembles interphase NPC assembly. Thus, it is plausible that retention of the basket is mediated through Nup60’s association with the segregating chromosomes and that disruption of the basket leaves the newly formed nuclei unable to properly repopulate their membranes with NPCs. It is unclear, however, why an effect on sporulation would only manifest when the cells are also *ndj1* or *csm4*.

The mechanism by which Nup60 promotes meiosis and sporulation once it has bound Nup2 is still an open question. Nup136, the functional homolog of Nup153 in *Arabidopsis*, helps remove meiotic interlocks by promoting chromosome movement (Martinez-Garcia *et al*. 2018). While *nup2* mutants do not show any obvious defect in rapid prophase movement (Chu et al, 2017), it is possible that Nup2/Nup60 influences chromosome movement in more subtle ways. Nup60 may also be acting indirectly via effects on gene expression. Nup60 represses transcription of *CLN2* by recruiting the gene to the nuclear envelope (Kumar *et al*. 2018). Interestingly, deacetylation of a lysine residue in the C-terminus of Nup60 prevents it from binding to the *CLN2* promoter. Alternatively, Nup2/Nup60 might be acting through Hop1 or Pch1. Nup2 facilitates removal of Hop1 from interstitial regions of chromosomes, allowing double strand break potential to persist near the chromosome ends and helping to ensure that short chromosomes form a crossover (Subramanian *et al*. 2019). Nup2 has also been shown to promote association of the checkpoint protein Pch2 with meiotic chromosomes and to control the distribution of Pch2 between the nucleus and the cytoplasm (Herruzo *et al*. 2021). It will be interesting to see if Nup2’s effect on either Hop1 or Pch2 requires Nup60. Finally, several landmarks of meiotic prophase are delayed in the *nup2* mutant, including homolog pairing, double-strand break repair, and synapsis, and this delay is much worse in an *ndj1* mutant background (Chu et al, 2017). On the other hand, double-strand breaks and the products of recombination (both crossovers and noncrossovers) are slightly elevated in the *nup2* mutant. It would not be surprising if *nup60* mutants exhibited these same phenotypes.

The NPC was initially defined by its role in nucleocytoplasmic trafficking (Wente and Rout 2010) but also helps to maintain the functional organization of chromatin within the nucleus. This organization influences a variety of DNA transactions including gene expression and DNA repair. Our work has uncovered a mechanism linking the NPC basket proteins Nup60 and Nup2 to sporulation, with Nup60 playing the major role and a 42-aa region of Nup2 acting to alter Nup60’s activity. Nup60’s C-terminus appears to inhibit the N-terminus, and binding of Nup2 relieves this inhibition. Whether the interaction between Nup2 and Nup60 is regulatory in nature and whether it is preserved in other processes involving Nup2/Nup60 binding to chromatin remains to be determined.

## Data availability

Strains and plasmids are available upon request. All supplemental figures, tables, and files and raw image data will be deposited in the Dryad repository upon acceptance for publication; doi TBD. Supplemental information contains supplemental figures, tables, and methods; Supplemental File S1 contains sporulation data; File S2 contains fluorescence measurements for Figure 3AB; File S3 contains fluorescence measurements for Figure 3CD; File S4 contains fluorescence measurements for Figure 6; File S5 contains fluorescence measurements for Figure 8; File S6 contains time course data.

## Acknowledgements

We thank members of the Burgess lab for stimulating discussions and Carly Cheung and Daniel Chu for technical support with plasmids and yeast strains. We are grateful to Shannon Owens and Amy MacQueen for advice on the chromosome spread procedure. This work was funded by NIH R01 GM701195 to SMB.

## References

Aguilera P., J. Whalen, C. Minguet, D. Churikov, C. Freudenreich, et al., 2020 The nuclear pore complex prevents sister chromatid recombination during replicative senescence. Nat. Commun. 11: 160.

Arora C., K. Kee, S. Maleki, and S. Keeney, 2004 Antiviral protein Ski8 is a direct partner of Spo11 in meiotic DNA break formation, independent of its cytoplasmic role in RNA metabolism. Mol. Cell 13: 549–559.

Bermejo R., T. Capra, R. Jossen, A. Colosio, C. Frattini, et al., 2011 The replication checkpoint protects fork stability by releasing transcribed genes from nuclear pores. Cell 146: 233–246.

Boeke J. D., F. La Croute, and G. R. Fink, 1984 A positive selection for mutants lacking orotidine-5′-phosphate decarboxylase activity in yeast: 5-fluoro-orotic acid resistance. Mol. Gen. Genet. 197: 345–346.

Brickner D. G., S. Ahmed, L. Meldi, A. Thompson, W. Light, et al., 2012 Transcription factor binding to a DNA zip code controls interchromosomal clustering at the nuclear periphery. Dev. Cell 22: 1234–1246.

Brickner D. G., C. Randise-Hinchliff, M. Lebrun Corbin, J. M. Liang, S. Kim, et al., 2019 The Role of Transcription Factors and Nuclear Pore Proteins in Controlling the Spatial Organization of the Yeast Genome. Dev. Cell 49: 936–947.e4.

Buchwalter A. L., Y. Liang, and M. W. Hetzer, 2014 Nup50 is required for cell differentiation and exhibits transcription-dependent dynamics. Mol. Biol. Cell 25: 2472–2484.

Burke B., 2018 LINC complexes as regulators of meiosis. Current Opinion in Cell Biology 52: 22–29.

Casolari J. M., C. R. Brown, S. Komili, J. West, H. Hieronymus, et al., 2004 Genome-wide localization of the nuclear transport machinery couples transcriptional status and nuclear organization. Cell 117: 427–439.

Chu D. B., T. Gromova, T. A. C. Newman, and S. M. Burgess, 2017 The Nucleoporin Nup2 Contains a Meiotic-Autonomous Region that Promotes the Dynamic Chromosome Events of Meiosis. Genetics 206: 1319–1337.

Conrad M. N., C.-Y. Lee, G. Chao, M. Shinohara, H. Kosaka, et al., 2008 Rapid telomere movement in meiotic prophase is promoted by NDJ1, MPS3, and CSM4 and is modulated by recombination. Cell 133: 1175–1187.

Denning D., B. Mykytka, N. P. Allen, L. Huang, Al Burlingame, et al., 2001 The nucleoporin Nup60p functions as a Gsp1p-GTP-sensitive tether for Nup2p at the nuclear pore complex. J. Cell Biol. 154: 937–950.

Denning D. P., S. S. Patel, V. Uversky, A. L. Fink, and M. Rexach, 2003 Disorder in the nuclear pore complex: the FG repeat regions of nucleoporins are natively unfolded. Proc. Natl. Acad. Sci. U. S. A. 100: 2450–2455.

Dilworth D. J., A. Suprapto, J. C. Padovan, B. T. Chait, R. W. Wozniak, et al., 2001 Nup2p dynamically associates with the distal regions of the yeast nuclear pore complex. J. Cell Biol. 153: 1465–1478.

Dilworth D. J., A. J. Tackett, R. S. Rogers, E. C. Yi, R. H. Christmas, et al., 2005 The mobile nucleoporin Nup2p and chromatin-bound Prp20p function in endogenous NPC-mediated transcriptional control. J. Cell Biol. 171: 955–965.

Edelstein A., N. Amodaj, K. Hoover, R. Vale, and N. Stuurman, 2010 Computer control of microscopes using µManager. Curr. Protoc. Mol. Biol. Chapter 14: Unit14.20.

Fuchs J., and J. Loidl, 2004 Behaviour of nucleolus organizing regions (NORs) and nucleoli during mitotic and meiotic divisions in budding yeast. Chromosome Res. 12: 427–438.

Grubb J., M. S. Brown, and D. K. Bishop, 2015 Surface Spreading and Immunostaining of Yeast Chromosomes. J. Vis. Exp. e53081.

Hase M. E., and V. C. Cordes, 2003 Direct interaction with nup153 mediates binding of Tpr to the periphery of the nuclear pore complex. Mol. Biol. Cell 14: 1923–1940.

Herruzo E., A. Lago-Maciel, S. Baztán, B. Santos, J. A. Carballo, et al., 2021 Pch2 orchestrates the meiotic recombination checkpoint from the cytoplasm. PLoS Genet. 17: e1009560.

Hood J. K., J. M. Casolari, and P. A. Silver, 2000 Nup2p is located on the nuclear side of the nuclear pore complex and coordinates Srp1p/importin-alpha export. J. Cell Sci. 113 (Pt 8): 1471–1480.

Ishii K., G. Arib, C. Lin, G. Van Houwe, and U. K. Laemmli, 2002 Chromatin Boundaries in Budding Yeast. Cell 109: 551–562.

Jumper J., R. Evans, A. Pritzel, T. Green, M. Figurnov, et al., 2021 Highly accurate protein structure prediction with AlphaFold. Nature 596: 583–589.

Kalverda B., H. Pickersgill, V. V. Shloma, and M. Fornerod, 2010 Nucleoporins directly stimulate expression of developmental and cell-cycle genes inside the nucleoplasm. Cell 140: 360–371.

Kim S., I. Liachko, D. G. Brickner, K. Cook, W. S. Noble, et al., 2017 The dynamic three-dimensional organization of the diploid yeast genome. Elife 6. https://doi.org/10.7554/eLife.23623

King G. A., J. S. Goodman, J. G. Schick, K. Chetlapalli, D. M. Jorgens, et al., 2019 Meiotic cellular rejuvenation is coupled to nuclear remodeling in budding yeast. Elife 8. https://doi.org/10.7554/eLife.47156

Kumar A., P. Sharma, M. Gomar-Alba, Z. Shcheprova, A. Daulny, et al., 2018 Daughter-cell-specific modulation of nuclear pore complexes controls cell cycle entry during asymmetric division. Nat. Cell Biol. 20: 432–442.

Longtine M. S., A. McKenzie 3rd, D. J. Demarini, N. G. Shah, A. Wach, et al., 1998 Additional modules for versatile and economical PCR-based gene deletion and modification in Saccharomyces cerevisiae. Yeast 14: 953–961.

Lui D., and S. M. Burgess, 2009 Measurement of Spatial Proximity and Accessibility of Chromosomal Loci in Saccharomyces cerevisiae Using Cre /loxP Site-Specific Recombination, pp. 55–63 in Meiosis: Volume 1, Molecular and Genetic Methods, edited by Keeney S. Humana Press, Totowa, NJ.

Makise M., D. R. Mackay, S. Elgort, S. S. Shankaran, S. A. Adam, et al., 2012 The Nup153-Nup50 Protein Interface and Its Role in Nuclear Import. Journal of Biological Chemistry 287: 38515–38522.

Markossian S., S. Suresh, A. H. Osmani, and S. A. Osmani, 2015 Nup2 requires a highly divergent partner, NupA, to fulfill functions at nuclear pore complexes and the mitotic chromatin region. Mol. Biol. Cell 26: 605–621.

Martinez-Garcia M., V. Schubert, K. Osman, A. Darbyshire, E. Sanchez-Moran, et al., 2018 TOPII and chromosome movement help remove interlocks between entangled chromosomes during meiosis. J. Cell Biol. 217: 4070–4079.

McCloy R. A., S. Rogers, C. E. Caldon, T. Lorca, A. Castro, et al., 2014 Partial inhibition of Cdk1 in G 2 phase overrides the SAC and decouples mitotic events. Cell Cycle 13: 1400–1412.

McKee A. H., and N. Kleckner, 1997 A general method for identifying recessive diploid-specific mutations in Saccharomyces cerevisiae, its application to the isolation of mutants blocked at intermediate stages of meiotic prophase and characterization of a new gene SAE2. Genetics 146: 797–816.

Mehla J., J. H. Caufield, N. Sakhawalkar, and P. Uetz, 2017 A comparison of two-hybrid approaches for detecting protein-protein interactions. Methods Enzymol. 586: 333–358.

Mészáros N., J. Cibulka, M. J. Mendiburo, A. Romanauska, M. Schneider, et al., 2015 Nuclear pore basket proteins are tethered to the nuclear envelope and can regulate membrane curvature. Dev. Cell 33: 285–298.

Pyhtila B., and M. Rexach, 2003 A gradient of affinity for the karyopherin Kap95p along the yeast nuclear pore complex. J. Biol. Chem. 278: 42699–42709.

Rabut G., V. Doye, and J. Ellenberg, 2004 Mapping the dynamic organization of the nuclear pore complex inside single living cells. Nat. Cell Biol. 6: 1114–1121.

Raices M., and M. A. D’Angelo, 2021 Structure, Maintenance, and Regulation of Nuclear Pore Complexes: The Gatekeepers of the Eukaryotic Genome. Cold Spring Harb. Perspect. Biol. https://doi.org/10.1101/cshperspect.a040691

Rockmill B., 2009 Chromosome spreading and immunofluorescence methods in Saccharomyes cerevisiae. Methods Mol. Biol. 558: 3–13.

Scherthan H., H. Wang, C. Adelfalk, E. J. White, C. Cowan, et al., 2007 Chromosome mobility during meiotic prophase in Saccharomyces cerevisiae. Proc. Natl. Acad. Sci. U. S. A. 104: 16934–16939.

Schindelin J., I. Arganda-Carreras, E. Frise, V. Kaynig, M. Longair, et al., 2012 Fiji: an open-source platform for biological-image analysis. Nat. Methods 9: 676–682.

Schmid M., G. Arib, C. Laemmli, J. Nishikawa, T. Durussel, et al., 2006 Nup-PI: the nucleopore-promoter interaction of genes in yeast. Mol. Cell 21: 379–391.

Schwartz M., A. Travesa, S. W. Martell, and D. J. Forbes, 2015 Analysis of the initiation of nuclear pore assembly by ectopically targeting nucleoporins to chromatin. Nucleus 6: 40–54.

Sheff M. A., and K. S. Thorn, 2004 Optimized cassettes for fluorescent protein tagging inSaccharomyces cerevisiae. Yeast 21: 661–670.

Sistla S., J. V. Pang, C. X. Wang, and D. Balasundaram, 2007 Multiple conserved domains of the nucleoporin Nup124p and its orthologs Nup1p and Nup153 are critical for nuclear import and activity of the fission yeast Tf1 retrotransposon. Mol. Biol. Cell 18: 3692–3708.

Solsbacher J., P. Maurer, F. Vogel, and G. Schlenstedt, 2000 Nup2p, a yeast nucleoporin, functions in bidirectional transport of importin alpha. Mol. Cell. Biol. 20: 8468–8479.

Subramanian V. V., X. Zhu, T. E. Markowitz, L. A. Vale-Silva, P. A. San-Segundo, et al., 2019 Persistent DNA-break potential near telomeres increases initiation of meiotic recombination on short chromosomes. Nat. Commun. 10: 970.

Suresh S., S. Markossian, A. H. Osmani, and S. A. Osmani, 2017 Mitotic nuclear pore complex segregation involves Nup2 in Aspergillus nidulans. J. Cell Biol. 216: 2813–2826.

Suresh S., S. Markossian, A. H. Osmani, and S. A. Osmani, 2018 Nup2 performs diverse interphase functions in Aspergillus nidulans. Mol. Biol. Cell 29: 3144–3154.

Vollmer B., M. Lorenz, D. Moreno-Andrés, M. Bodenhöfer, P. De Magistris, et al., 2015 Nup153 Recruits the Nup107-160 Complex to the Inner Nuclear Membrane for Interphasic Nuclear Pore Complex Assembly. Dev. Cell 33: 717–728.

Wanat J. J., K. P. Kim, R. Koszul, S. Zanders, B. Weiner, et al., 2008 Csm4, in collaboration with Ndj1, mediates telomere-led chromosome dynamics and recombination during yeast meiosis. PLoS Genet. 4: e1000188.

Wente S. R., and M. P. Rout, 2010 The nuclear pore complex and nuclear transport. Cold Spring Harb. Perspect. Biol. 2: a000562.

Zhao X., C.-Y. Wu, and G. Blobel, 2004 Mlp-dependent anchorage and stabilization of a desumoylating enzyme is required to prevent clonal lethality. J. Cell Biol. 167: 605–611.

