## Supplemental information for "The Nup2 meiotic-autonomous region relieves autoinhibition of Nup60 to promote progression of meiosis and sporulation in *Saccharomyces cerevisiae*"

**Supplemental Figures, Tables, Methods, and References for Komachi and Burgess, 2021**

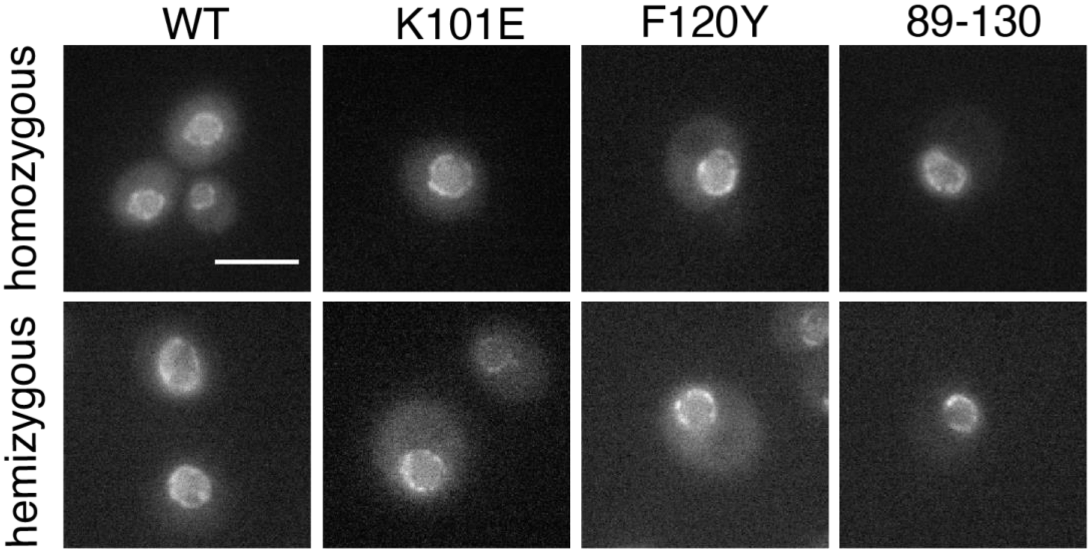

**Figure S1.** Effect of hemizygosity on localization of mar(K101E), mar(F120Y), and mar(89-130) GFP fusions. Top panels show fluorescent images of the homozygous strains, bottom panels show hemizygous strains. The MAR allele depicted is indicated at the top of each column. The scale bar in the wild-type MAR-GFP panel represents 5 μm.

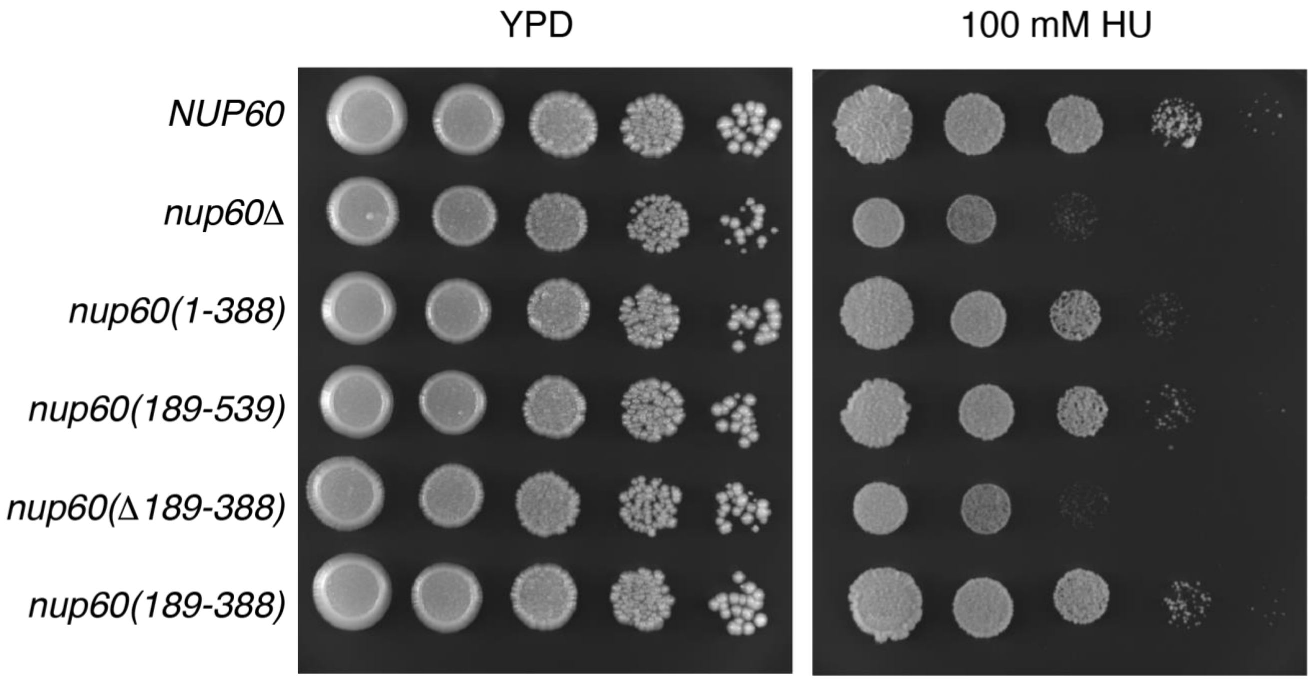

**Figure S2.** Complementation of HU sensitivity by *NUP60* fragments. Cells were grown to mid-log phase in liquid YPD and diluted to an $OD_{600}$ of 1. Serial 10-fold dilutions were spotted onto YPD (left) or YPD plus 100 mM hydroxyurea (right) and grown at 30˚C for 48 hours and 96 hours, respectively. The strains are isogenic to the wild-type (*NUP60*) strain: *NUP60* (SBY1903), *nup60Δ* (SBY5217), *nup60(1-388)* (SBY6423), *nup60(189-539)* (SBY6426), *nup60(Δ189-388)* (SBY6429), *nup60(189-388)* (SBY6420).

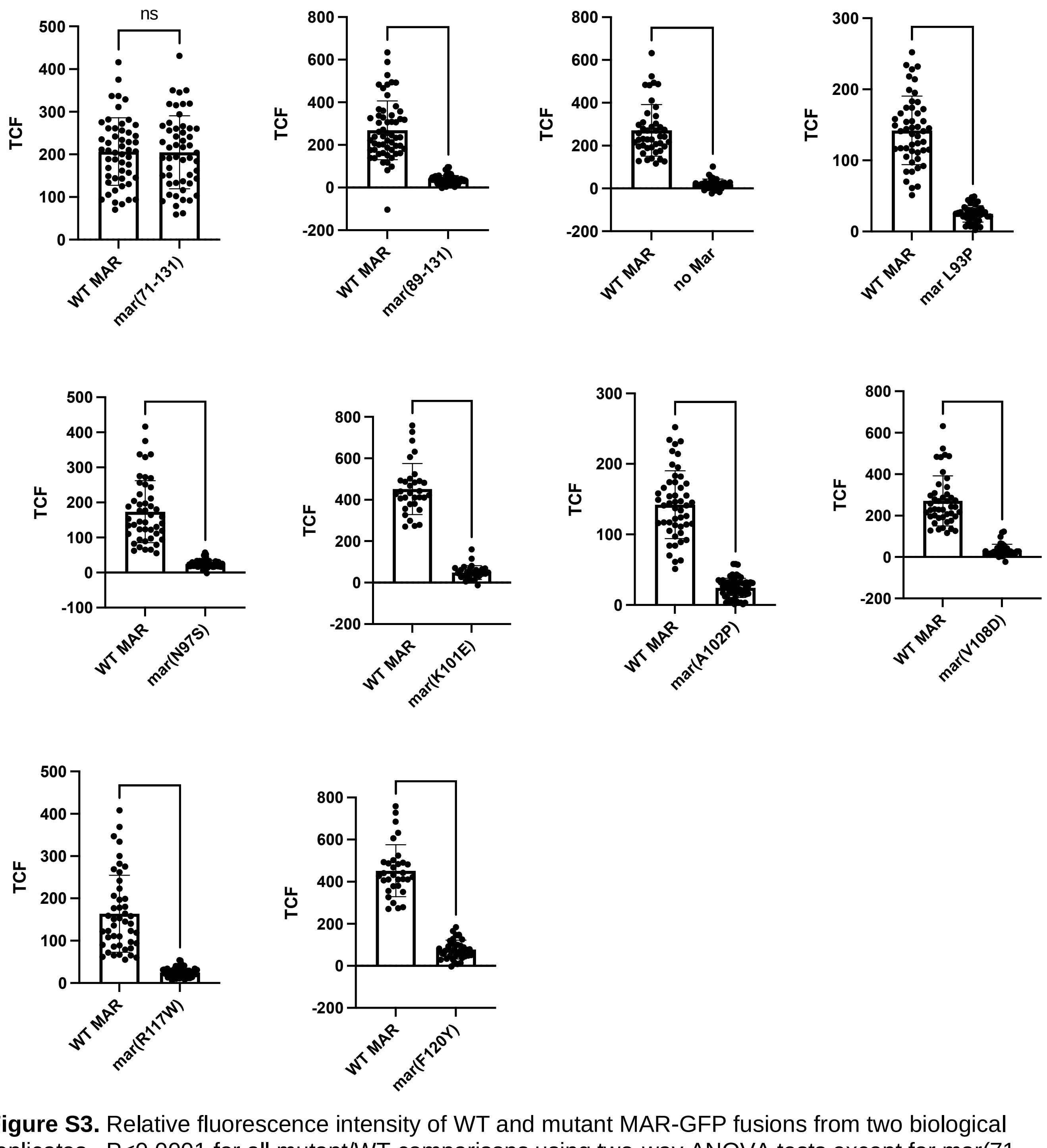

**Figure S3.** Relative fluorescence intensity of WT and mutant MAR-GFP fusions from two biological replicates. P<0.0001 for all mutant/WT comparisons using two-way ANOVA tests except for mar(71-130) in which P=0.95.

**A**

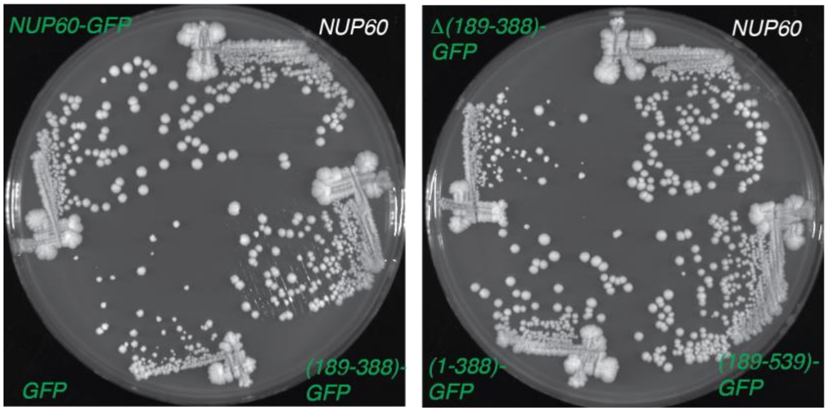

**B**

| Relevant genotype | Growth | Sporulation efficiency |
|---|---|---|
| *NUP60-GFP* | normal | 72 ± 7 |
| *GFP* | poor | 15 ± 5 |
| *(189-388)-GFP* | normal | 20 ± 6 |
| *(1-388)-GFP* | normal | 68 ± 7 |
| *(189-539)-GFP* | normal | 15 ± 3 |
| *(Δ189-388)-GFP* | poor | 16 ± 4 |

**Figure S4.** Complementation of growth and sporulation by Nup60 truncations fused to GFP. (A) Yeast strains streaked out onto YPD plates. All strains are isogenic to the wild-type (*NUP60*) strain: *NUP60* (SBY1899), *NUP60-GFP* (SBY6304), *GFP* (SBY6307), *nup60(189-388)-GFP* (SBY6310), *nup60(1-388)* (SBY6313), *nup60(189-539)* (SBY6316), *nup60(Δ189-388)-GFP* (SBY6382). (B) Summary of growth and sporulation of strains expressing truncated Nup60-GFP fusions. The strains used to assay sporulation efficiency are in an *ndj1* background and are all isogenic to the *NUP60-GFP* strain: *NUP60-GFP* (SBY6320), *GFP* (SBY6323), *nup60(189-388)-GFP* (SBY6326), *nup60(1-388)* (SBY6329), *nup60(189-539)* (SBY6332), *nup60(Δ189-388)-GFP* (SBY6335).

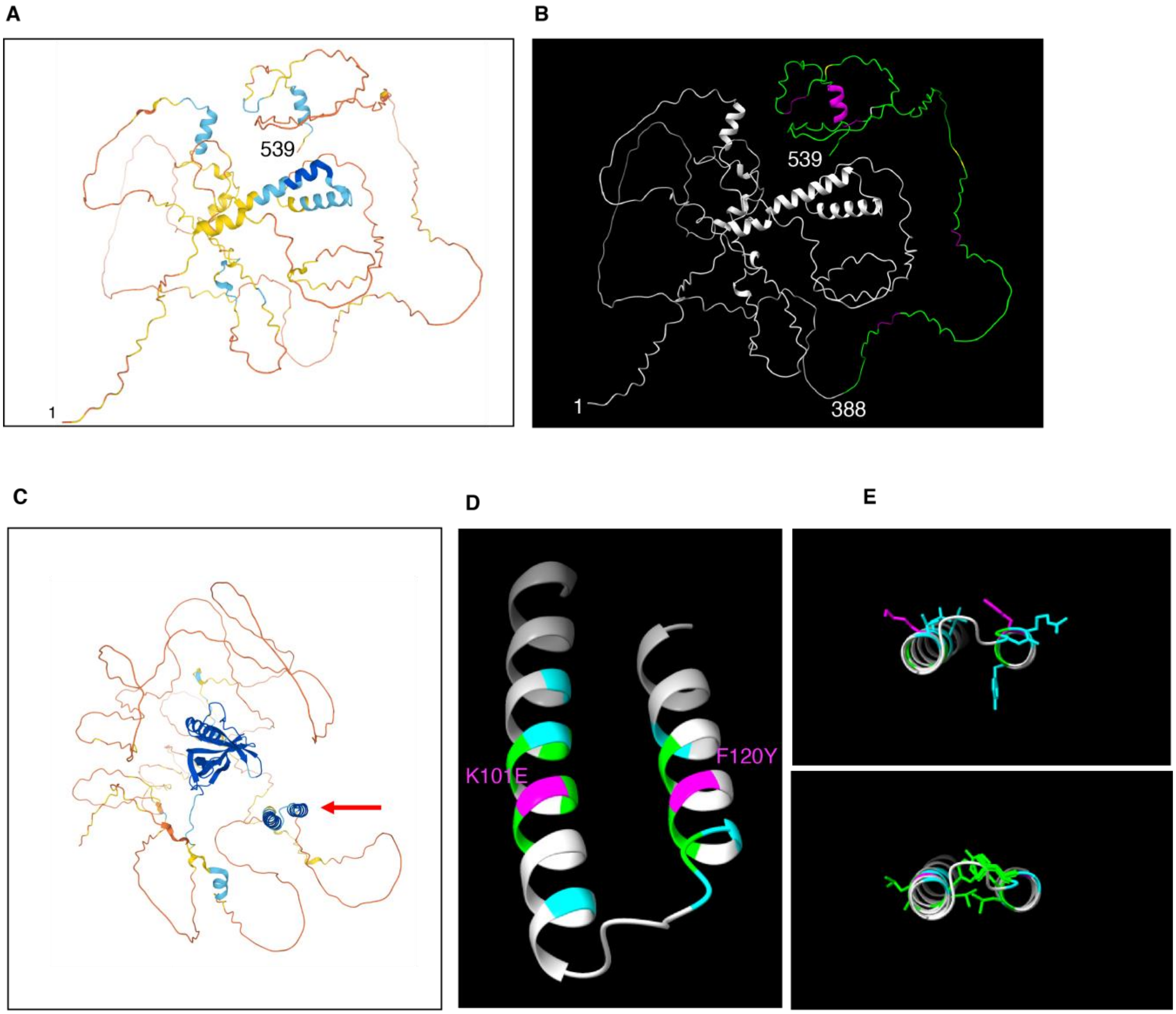

**Figure S5.** AlphaFold predictions of the structures of Nup60 and Nup2. (A) Nup60 predicted structure from the AlphaFold v2.0 database updated on 1 July 2021 (Jumper *et al.* 2021). Regions are color-coded according to AlphaFold confidence scores based on the local distance difference test (LDDT) (Mariani *et al.* 2013): Very high (pLDDT > 90); Confident (90 > pLDDT > 70); Low (70 > pLDDT > 50); Very low (pLDDT < 50). The first and last amino acids are numbered. (B) Nup60 predicted structure, with the region sufficient for sporulation function (aa 1-388) colored gray and the region containing the inhibitory domain (aa 389-539) colored green. A predicted helix near the extreme C-terminus is colored magenta; the FxF repeats (aa 399-401, 427-429, 469-471, 509-511), purple; SUMOylation sites (aa 440, 442, 505), yellow; a known acetylation site (aa 467), white. (C) AlphaFold v2.0 prediction of the structure of Nup2, with regions color-coded according to confidence scores described in (A). The MAR (indicated by the red arrow) and Ran GTPase binding domain are the only two regions where structure can be predicted with high confidence. (D) Blow-up of the helices that make up the MAR, rotated approximately 90˚ around the x-axis with respect to (C). The positions of the mutations are marked by different colors: mutations that do not disrupt localization to the nuclear envelope (K101 and F120) are colored magenta; mutations that disrupt localization but not binding to full-length Nup60 in Y2H, cyan; mutations that disrupt localization and all Y2H interaction, green. (E) Helices of the MAR in the same orientation as in (C), with side chains shown. The top panel highlights mutations that do not disrupt binding to full-length Nup60 in the Y2H assay; the bottom panel highlights mutations that disrupt Y2H to full-length Nup60. Images in B, D, and E were created using AlphaFold v2.0 coordinates and UCSF ChimeraX version 1.2.5 (Pettersen *et al.* 2021).

A

| Relevant genotype | Sporulation efficiency |
|---|---|
| *3HA-NUP60* | 82 ± 8 |
| *3HA-NUP60 ndj1* | 21 ± 5 |
| *3HA-NUP60 nup2* | 75 ± 5 |
| *3HA-NUP60 ndj1 nup2* | 14 ± 2 |

B

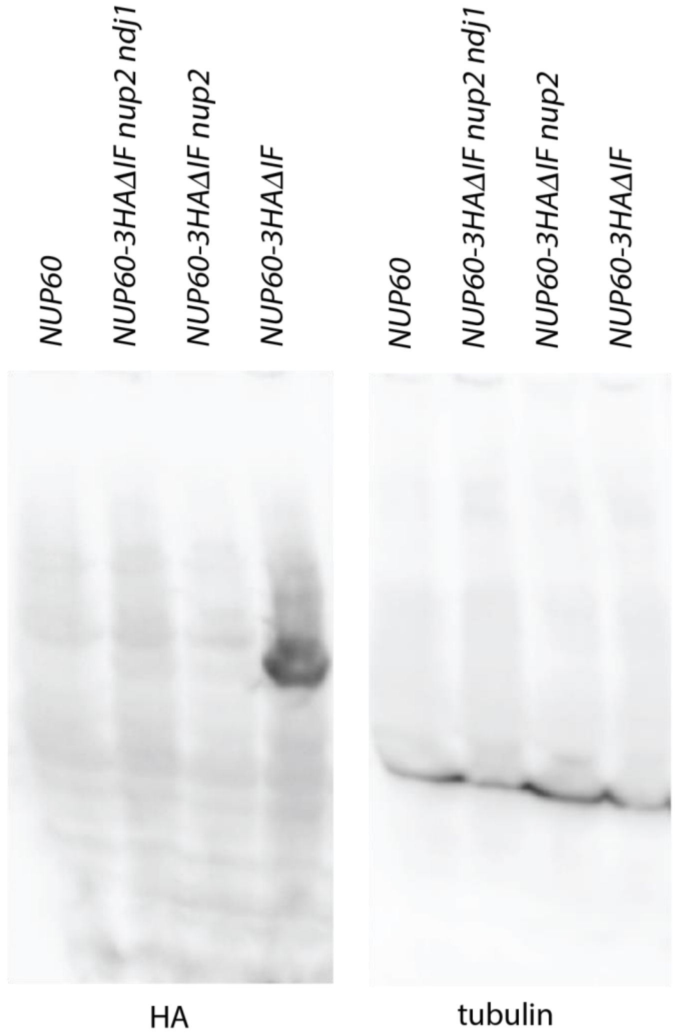

**Figure S6.** Disruption of normal Nup60 function by HA tagging at the N- or C-terminus. (A) Table showing the sporulation efficiency of *3HA-NUP60* strains. N-terminally tagged Nup60 is unable to promote sporulation in the absence of Ndj1. (B) Western blot of *NUP60* or *NUP60-3HA* strains probed with an antibody against HA (left) or β-tubulin (right). The linker between Nup60 and the HA tag has been modified to remove two amino acids known to destabilize HA tagged proteins in yeast (Saiz-Baggetto *et al.* 2017). The fusion is strongly expressed in wild-type cells but cannot be detected in extracts from *nup2* strains. Because *NUP60-3HA nup2* strains grow well, the fusion is probably being only partially degraded, leaving enough Nup60 to support cell growth. It is unclear why Nup60-3HA is only unstable in the absence of Nup2.

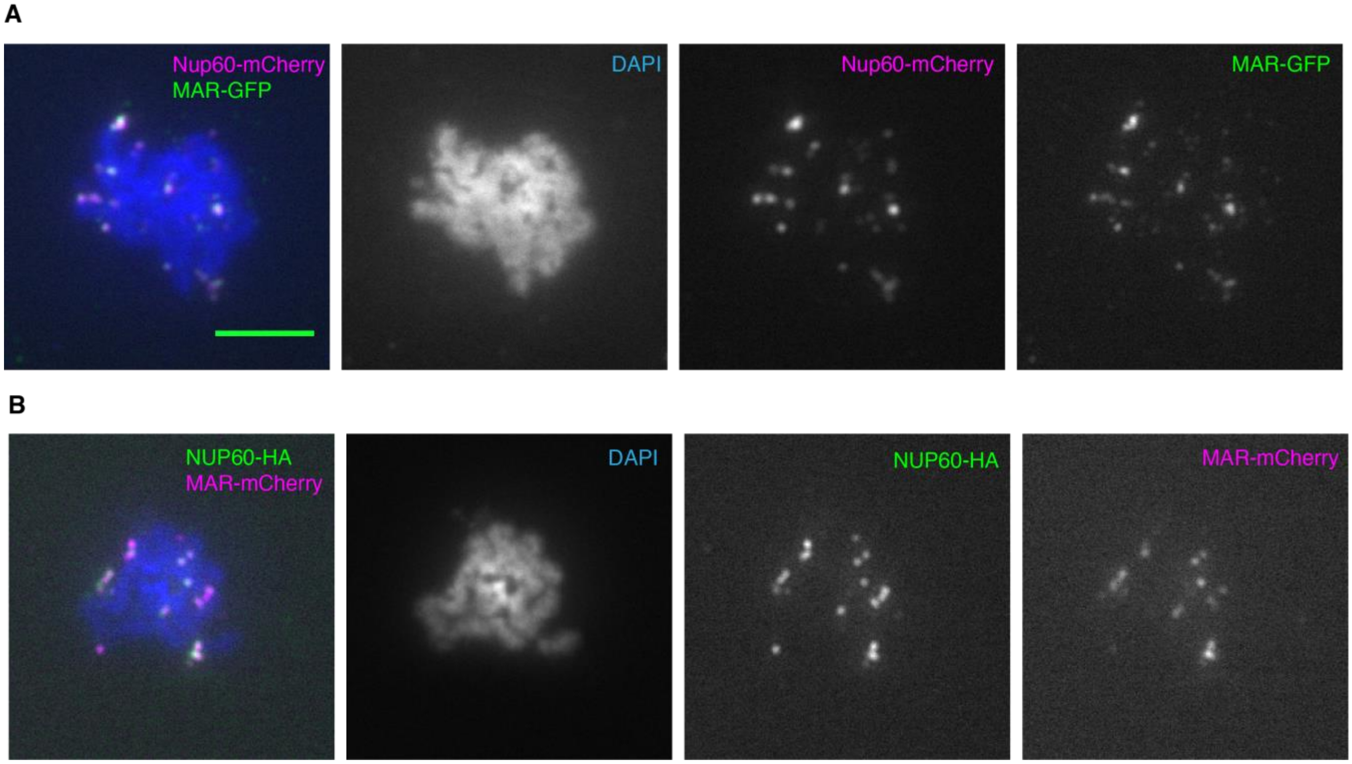

**Figure S7.** Binding of meiotic chromosomes by Nup60-mCherry and Nup60-3HA. (A) Representative meiotic chromosome spreads from a strain expressing Nup60-mCherry and MAR-GFP (SBY6481) . Left to right are the merged image and images from the DAPI, TRITC, and FITC channels. (B) Representative meiotic chromosome spreads from a strain expressing Nup60-3HA and MAR-mCherry (SBY6484). Left to right are the merged image and images from the DAPI, FITC, and TRITC channels. The GFP fusions were detected using a polyclonal antibody to GFP and an Alexa Fluor 488-conjugated secondary antibody. Nup60-mCherry was detected using a polyclonal antibody to mCherry and an Alexa Fluor 594-conjugated secondary. Nup60-3HA was detected using a polyclonal antibody to HA and an Alexa Fluor 594-conjugated secondary. The scale bar in the upper left panel represents 5 $\mu$m.

**Table S1.** Yeast strains used in this study

| Strain | Genotype | Background | Figure Panel | Reference/ |
|---|---|---|---|---|
| SBY6259 | *MATa ho::hisG leu2::hisG ura3::hisG GAL3(?) his4::LEU2-(NBam) nup2(Δ51-175) ndj1::TRP1* | SK1 | Figure 1B (Screen) | |
| SBY6260 | *MATalpha ho::hisG leu2::hisG ura3(ΔSma-PstI) HIS4::LEU2-(NBam) nup2::KanMx ndj1::KanMx* | SK1 | Figure 1B (Screen) | |
| SBY1899 | *MATalpha ho::hisG leu2::hisG ura3(ΔSma-Pst) his4-X::LEU2-(NBam)-URA3* | SK1 | Wild-type parent of SBY1903 | (Chu *et al.* 2017) |
| SBY1900 | *MATa ho::hisG leu2::hisG ura3(ΔSma-Pst) HIS4::LEU2-(NBam)* | SK1 | Wild-type parent of SBY1903 | (Chu *et al.* 2017) |
| SBY1903 | *MATa ho::hisG leu2::hisG ura3(ΔSma-Pst) HIS4::LEU2-(NBam)*<br>*MATalpha ho::hisG leu2::hisG ura3(ΔSma-Pst) his4-X::LEU2-(NBam)-URA3* | SK1 | Wild-type diploid strain | (Chu *et al.* 2017) |
| SBY6062 | *MATa ho::hisG leu2::hisG ura3(ΔSma-Pst) HIS4::LEU2-(NBam) ndj1::KanMx nup2(51-175)-GFP::CaURA3*<br>*MATalpha ho::hisG leu2::hisG ura3(ΔSma-Pst) his4-X::LEU2-(NBam) ndj1::KanMx nup2(51-175)-GFP::CaURA3* | SK1 | Table 1 (wild-type MAR) | |
| SBY6062* | *MATa ho::hisG leu2::hisG ura3(ΔSma-Pst) HIS4::LEU2-(NBam) ndj1::KanMx nup2(51-175)*-GFP::CaURA3*<br>*MATalpha ho::hisG leu2::hisG ura3(ΔSma-Pst) his4-X::LEU2-(NBam)-URA3 ndj1::KanMx nup2(51-175)*-GFP::CaURA3* | SK1 | Table 1 (Point mutant versions of SBY6062) | |
| SBY6059 | *MATa ho::hisG leu2::hisG ura3(ΔSma-Pst) HIS4::LEU2-(NBam) ndj1::KanMx nup2(Δ1-720)-GFP::CaURA3*<br>*MATalpha ho::hisG leu2::hisG ura3(ΔSma-Pst) his4-X::LEU2-(NBam)-URA3 ndj1::KanMx nup2(Δ1-720)-GFP::CaURA3* | SK1 | Table 1 (GFP control) | |
| SBY6065 | *MATa ho::hisG leu2::hisG ura3(ΔSma-Pst) HIS4::LEU2-(NBam) ndj1::KanMx nup2(71-130)-GFP::CaURA3* | SK1 | Table 1 | |

| | | | | |
|---|---|---|---|---|
| | *MATalpha ho::hisG leu2::hisG ura3(ΔSma-Pst) his4-X::LEU2-(NBam)-URA3 ndj1::KanMx nup2(71-175)-GFP::CaURA3* | | | |
| SBY6266 | *MATa ho::hisG leu2::hisG ura3(ΔSma-Pst) HIS4::LEU2-(NBam) ndj1::KanMx nup2(89-130)-GFP::CaURA3* | SK1 | Table 1 | |
| | *MATalpha ho::hisG leu2::hisG ura3(ΔSma-Pst) his4-X::LEU2-(NBam)-URA3 ndj1::KanMx nup2(89-175)-GFP::CaURA3* | | | |
| SBY6414 | *MATa ho::hisG leu2::hisG ura3(ΔSma-Pst) HIS4::LEU2-(NBam) ndj1::KanMx nup2(89-175)-GFP::CaURA3* | SK1 | Table 1 | |
| | *MATalpha ho::hisG leu2::hisG ura3(ΔSma-Pst) his4-X::LEU2-(NBam)-URA3 ndj1::KanMx nup2(89-175)-GFP::CaURA3* | | | |
| SBY3981 | *MATalpha ho::hisG leu2::hisG ura3(ΔSma-Pst) his4-X::LEU2-(NBam)-URA3 nup2:KanMx ndj1::KanMx* | SK1 | Table 1 | (Chu *et al.* 2017) |
| SBY6114 | *MATa ho::hisG leu2::hisG ura3(ΔSma-Pst) HIS4::LEU2-(NBam) nup2(51-175)*$^{K101E}$*-GFP::CaURA3 ndj1::KanMx* | SK1 | Table 1 | |
| SBY6081 | *MATa ho::hisG leu2::hisG ura3(ΔSma-Pst) HIS4::LEU2-(NBam) nup2(51-175)*$^{F120Y}$*-GFP::CaURA3 ndj1::KanMx* | SK1 | Table 1 | |
| SBY6264 | *MATa ho::hisG leu2::hisG ura3(ΔSma-Pst) HIS4::LEU2-(NBam) nup2(89-130)-GFP::CaURA3 ndj1::KanMx* | SK1 | Table 1 | |
| SBY6413 | *MATa ho::hisG leu2::hisG ura3(ΔSma-Pst) HIS4::LEU2-(NBam) nup2(89-175)-GFP::CaURA3 ndj1::KanMx* | SK1 | Table 1 | |
| SBY6068 | *MATa ho::hisG leu2::hisG ura3(ΔSma-Pst) HIS4::LEU2-(NBam) nup2(51-175)-GFP::CaURA3* | SK1 | Figure 1D (WT control) | |
| | *MATalpha ho::hisG leu2::hisG ura3(ΔSma-Pst) his4-X::LEU2-(NBam)-URA3 nup2(51-175)-GFP::CaURA3* | | | |
| SBY6068* | *MATa ho::hisG leu2::hisG ura3(ΔSma-Pst) HIS4::LEU2-(NBam) nup2(51-175)*-GFP::CaURA3* | SK1 | Figure 1D (point mutant versions of SBY6068) | |

| | | | | |
|---|---|---|---|---|
| | *MATalpha ho::hisG leu2::hisG ura3(ΔSma-Pst) his4-X::LEU2-(NBam)-URA3 nup2(51-175)*-GFP::CaURA3* | | | |
| SBY6074 | *MATa ho::hisG leu2::hisG ura3(ΔSma-Pst) HIS4::LEU2-(NBam) nup2(Δ1-720)-GFP::CaURA3*<br>*MATalpha ho::hisG leu2::hisG ura3(ΔSma-Pst) his4-X::LEU2-(NBam)-URA3 nup2(Δ1-720)-GFP::CaURA3* | SK1 | Figure 1D (GFP control) | |
| SBY6071 | *MATa ho::hisG leu2::hisG ura3(ΔSma-Pst) HIS4::LEU2-(NBam) nup2(71-130)-GFP::CaURA3*<br>*MATalpha ho::hisG leu2::hisG ura3(ΔSma-Pst) his4-X::LEU2-(NBam)-URA3 nup2(71-130)-GFP::CaURA3* | SK1 | Figure 1D | |
| SBY6263 | *MATa ho::hisG leu2::hisG ura3(ΔSma-Pst) HIS4::LEU2-(NBam) nup2(89-130)-GFP::CaURA3*<br>*MATalpha ho::hisG leu2::hisG ura3(ΔSma-Pst) his4-X::LEU2-(NBam)-URA3 nup2(89-130)-GFP::CaURA3* | SK1 | Figure 1D | |
| AH109 | *MATa, trp1–901, leu2–3, 112, ura3–52, his3–200, gal4Δ, gal80Δ, LYS2:: GAL1UAS-GAL1TATA-HIS3, GAL2UAS-GAL2TATA-ADE2, URA3:: MEL1UAS-MEL1TATA-lacZ.* | | Figure 2 | Clontech |
| SBY6054 | *MATa ho::hisG leu2::hisG ura3(ΔSma-Pst) HIS4::LEU2-(NBam) nup2(51-175)-GFP::CaURA3 ndt80::Hph*<br>*MATalpha ho::hisG leu2::hisG ura3(ΔSma-Pst) his4-X::LEU2-(NBam)-URA3 nup2(51-175)-GFP::CaURA3 ndt80::Hph* | SK1 | Figure 3A, B (wild-type control) | |
| SBY6119 | *MATa ho::hisG leu2::hisG ura3(ΔSma-Pst) HIS4::LEU2-(NBam) nup2(51-175)$^{K101E}$-GFP::CaURA3 ndt80::Hph*<br>*MATalpha ho::hisG leu2::hisG ura3(ΔSma-Pst) his4-X::LEU2-(NBam)-URA3 nup2(51-175)$^{K101E}$-GFP::CaURA3 ndt80::Hph* | SK1 | Figure 3A, B (K101E version of SBY6054) | |
| SBY6107 | *MATa ho::hisG leu2::hisG ura3(ΔSma-Pst) HIS4::LEU2-* | SK1 | Figure 3A, B (F120Y | |

| | | | |
|---|---|---|---|
| | *(NBam) nup2(51-175)*$^{F120Y}$*-GFP::CaURA3 ndt80::Hph*<br>*MATalpha ho::hisG leu2::hisG ura3(ΔSma-Pst) his4-X::LEU2-(NBam)-URA3 nup2(51-175)*$^{F120Y}$*-GFP::CaURA3 ndt80::Hph* | | version of SBY6054) |
| SBY6496 | *MATa ho::hisG leu2::hisG ura3(ΔSma-Pst) HIS4::LEU2-(NBam) nup2(51-175)*$^{N97S}$*-GFP::CaURA3 ndt80::Hph*<br>*MATalpha ho::hisG leu2::hisG ura3(ΔSma-Pst) his4-X::LEU2-(NBam)-URA3 nup2(51-175)*$^{N97S}$*-GFP::CaURA3 ndt80::Hph* | SK1 | Figure 3A, B (N97S version of SBY6054) |
| SBY6499 | *MATa ho::hisG leu2::hisG ura3(ΔSma-Pst) HIS4::LEU2-(NBam) nup2(51-175)*$^{V108D}$*-GFP::CaURA3 ndt80::Hph*<br>*MATalpha ho::hisG leu2::hisG ura3(ΔSma-Pst) his4-X::LEU2-(NBam)-URA3 nup2(51-175)*$^{V108D}$*-GFP::CaURA3 ndt80::Hph* | SK1 | Figure 3A, B (V108D version of SBY6054) |
| SBY6502 | *MATa ho::hisG leu2::hisG ura3(ΔSma-Pst) HIS4::LEU2-(NBam) nup2(51-175)*$^{R117W}$*-GFP::CaURA3 ndt80::Hph*<br>*MATalpha ho::hisG leu2::hisG ura3(ΔSma-Pst) his4-X::LEU2-(NBam)-URA3 nup2(51-175)*$^{R117W}$*-GFP::CaURA3 ndt80::Hph* | SK1 | Figure 3A, B (R117W version of SBY6054) |
| SBY6490 | *MATa ho::hisG leu2::hisG ura3(ΔSma-Pst) HIS4::LEU2-(NBam) nup2(51-175)*$^{L93P}$*-GFP::CaURA3 ndt80::Hph*<br>*MATalpha ho::hisG leu2::hisG ura3(ΔSma-Pst) his4-X::LEU2-(NBam)-URA3 nup2(51-175)*$^{L93P}$*-GFP::CaURA3 ndt80::Hph* | SK1 | Figure 3A, B (L93P version of SBY6054) |
| SBY6493 | *MATa ho::hisG leu2::hisG ura3(ΔSma-Pst) HIS4::LEU2-(NBam) nup2(51-175)*$^{A102P}$*-GFP::CaURA3 ndt80::Hph*<br>*MATalpha ho::hisG leu2::hisG ura3(ΔSma-Pst) his4-X::LEU2-(NBam)-URA3 nup2(51-175)*$^{A102P}$*-GFP::CaURA3 ndt80::Hph* | SK1 | Figure 3A, B (A102P version of SBY6054) |
| SBY6101 | *MATa ho::hisG leu2::hisG ura3(ΔSma-Pst) HIS4::LEU2-(NBam) nup2(Δ1-720)-GFP::CaURA3 ndt80::Hph* | SK1 | Figure 3A, B (GFP control) |

| | | | | |
|---|---|---|---|---|
| | *MATalpha ho::hisG leu2::hisG ura3(ΔSma-Pst) his4-X::LEU2-(NBam)-URA3 nup2(Δ1-720)-GFP::CaURA3 ndt80::Hph* | | | |
| SBY6269 | *MATa ho::hisG leu2::hisG ura3(ΔSma-Pst) HIS4::LEU2-(NBam) nup2(71-130)-GFP::CaURA3 ndt80::Hph*<br>*MATalpha ho::hisG leu2::hisG ura3(ΔSma-Pst) his4-X::LEU2-(NBam)-URA3 nup2(71-130)-GFP::CaURA3 ndt80::Hph* | SK1 | Figure 3A, B (mar(71-130) version of SBY6054) | |
| SBY6272 | *MATa ho::hisG leu2::hisG ura3(ΔSma-Pst) HIS4::LEU2-(NBam) nup2(89-130)-GFP::CaURA3 ndt80::Hph*<br>*MATalpha ho::hisG leu2::hisG ura3(ΔSma-Pst) his4-X::LEU2-(NBam)-URA3 nup2(89-130)-GFP::CaURA3 ndt80::Hph* | SK1 | Figure 3A, B (mar(89-130) version of SBY6054) | |
| SBY6278 | *MATa ho::hisG leu2::hisG ura3(ΔSma-Pst) HIS4::LEU2-(NBam) nup2(51-175)-GFP::CaURA3 ndt80::Hph nup60::KanMx*<br>*MATalpha ho::hisG leu2::hisG ura3(ΔSma-Pst) his4-X::LEU2-(NBam)-URA3 nup2(51-175)-GFP::CaURA3 ndt80::Hph nup60::KanMx* | SK1 | Figure 3C (nup60Δ version of SBY6054) | |
| SBY5213 | *MATalpha ho::hisG leu2::hisG ura3(ΔSma-Pst) his4-X::LEU2-(NBam)-URA3 ndj1::KanMx nup60::Hph* | SK1 | Figure 4B, D (growth) | |
| SBY6280 | *MATalpha ho::hisG leu2::hisG ura3(ΔSma-Pst) his4-X::LEU2-(NBam)-URA3 ndj1::KanMx nup60(189-388)* | SK1 | Figure 4B, D (growth) | |
| SBY6283 | *MATalpha ho::hisG leu2::hisG ura3(ΔSma-Pst) his4-X::LEU2-(NBam) ndj1::KanMx nup60(1-388)* | SK1 | Figure 4B, D (growth) | |
| SBY6286 | *MATalpha ho::hisG leu2::hisG ura3(ΔSma-Pst) his4-X::LEU2-(NBam)-URA3 ndj1::KanMx nup60(189-539)* | SK1 | Figure 4B, D (growth) | |
| SBY6302 | *MATalpha ho::hisG leu2::hisG ura3(ΔSma-Pst) his4-X::LEU2-(NBam)-URA3 ndj1::KanMx nup60(Δ189-388)* | SK1 | Figure 4B, D (growth) | |
| SBY5217 | *MATa ho::hisG leu2::hisG ura3(ΔSma-Pst) HIS4::LEU2-(NBam) nup60::Hph* | SK1 | Figures S2, 4B (HU | (Chu *et al.* 2017) different |

| | | | | |
|---|---|---|---|---|
| | *MATalpha ho::hisG leu2::hisG ura3(ΔSma-Pst) his4-X::LEU2-(NBam)-URA3 nup60::Hph* | | sensitivity, sporulation) | isolate of SBY5216 |
| SBY6420 | *MATa ho::hisG leu2::hisG ura3(ΔSma-Pst) HIS4::LEU2-(NBam) nup60(189-388)*<br>*MATalpha ho::hisG leu2::hisG ura3(ΔSma-Pst) his4-X::LEU2-(NBam)-URA3 nup60(189-388)* | SK1 | Figures S2, 4B (HU sensitivity, sporulation) | |
| SBY6423 | *MATa ho::hisG leu2::hisG ura3(ΔSma-Pst) HIS4::LEU2-(NBam) nup60(1-388)*<br>*MATalpha ho::hisG leu2::hisG ura3(ΔSma-Pst) his4-X::LEU2-(NBam)-URA3 nup60(1-388)* | SK1 | Figures S2, 4B (HU sensitivity, sporulation) | |
| SBY6426 | *MATa ho::hisG leu2::hisG ura3(ΔSma-Pst) HIS4::LEU2-(NBam) nup60(189-539)*<br>*MATalpha ho::hisG leu2::hisG ura3(ΔSma-Pst) his4-X::LEU2-(NBam)-URA3 nup60(189-539)* | SK1 | Figures S2, 4B (HU sensitivity, sporulation) | |
| SBY6429 | *MATa ho::hisG leu2::hisG ura3(ΔSma-Pst) HIS4::LEU2-(NBam) nup60(Δ189-388)*<br>*MATalpha ho::hisG leu2::hisG ura3(ΔSma-Pst) his4-X::LEU2-(NBam)-URA3 nup60(Δ189-388)* | SK1 | Figures S2, 4B (HU sensitivity, sporulation) | |
| SBY1904 | *MATa ho::hisG leu2::hisG ura3(ΔSma-Pst) HIS4::LEU2-(NBam) ndj1::KanMx*<br>*MATalpha ho::hisG leu2::hisG ura3(ΔSma-Pst) his4-X::LEU2-(NBam)-URA3 ndj1::KanMx* | SK1 | Figure 4B (sporulation) | (Chu *et al.* 2017) |
| SBY5222 | *MATa ho::hisG leu2::hisG ura3(ΔSma-Pst) HIS4::LEU2-(NBam) ndj1::KanMx nup60::Hph*<br>*MATalpha ho::hisG leu2::hisG ura3(ΔSma-Pst) his4-X::LEU2-(NBam)-URA3 ndj1::KanMx nup60::Hph* | SK1 | Figure 4B (sporulation) | |
| SBY6293 | *MATa ho::hisG leu2::hisG ura3(ΔSma-Pst) HIS4::LEU2-(NBam) ndj1::KanMx nup60(189-388)*<br>*MATalpha ho::hisG leu2::hisG ura3(ΔSma-Pst) his4-X::LEU2-(NBam)-URA3 ndj1::KanMx nup60(189-388)* | SK1 | Figures 4B, 5A-D (sporulation, meiotic time courses) | |
| SBY6296 | *MATa ho::hisG leu2::hisG ura3(ΔSma-Pst) HIS4::LEU2-(NBam) ndj1::KanMx nup60(1-388)* | SK1 | Figure 4B (sporulation) | |

| | | | |
|---|---|---|---|
| | *MATalpha ho::hisG leu2::hisG ura3(ΔSma-Pst) his4-X::LEU2-(NBam)-URA3 ndj1::KanMx nup60(1-388)* | | |
| SBY6299 | *MATa ho::hisG leu2::hisG ura3(ΔSma-Pst) HIS4::LEU2-(NBam) ndj1::KanMx nup60(189-539)*<br>*MATalpha ho::hisG leu2::hisG ura3(ΔSma-Pst) his4-X::LEU2-(NBam)-URA3 ndj1::KanMx nup60(189-539)* | SK1 | Figure 4B (sporulation) |
| SBY6302 | *MATa ho::hisG leu2::hisG ura3(ΔSma-Pst) HIS4::LEU2-(NBam) ndj1::KanMx nup60(Δ189-388)*<br>*MATalpha ho::hisG leu2::hisG ura3(ΔSma-Pst) his4-X::LEU2-(NBam)-URA3 ndj1::KanMx nup60(Δ189-388)* | SK1 | Figure 4B (sporulation) |
| SBY6304 | *MATa ho::hisG leu2::hisG ura3(ΔSma-Pst) HIS4::LEU2-(NBam) NUP60-GFP::KanMx*<br>*MATalpha ho::hisG leu2::hisG ura3(ΔSma-Pst) his4-X::LEU2-(NBam)-URA3 NUP60-GFP::KanMx* | SK1 | Figure S3A (GFP fusion growth) |
| SBY6307 | *MATa ho::hisG leu2::hisG ura3(ΔSma-Pst) HIS4::LEU2-(NBam) nup60(Δ1-539)-GFP::KanMx*<br>*MATalpha ho::hisG leu2::hisG ura3(ΔSma-Pst) his4-X::LEU2-(NBam)-URA3 nup60(Δ1-539)-GFP::KanMx* | SK1 | Figure S3A (GFP fusion growth) |
| SBY6310 | *MATa ho::hisG leu2::hisG ura3(ΔSma-Pst) HIS4::LEU2-(NBam) nup60(189-388)-GFP::KanMx*<br>*MATalpha ho::hisG leu2::hisG ura3(ΔSma-Pst) his4-X::LEU2-(NBam)-URA3 nup60(189-388)-GFP::KanMx* | SK1 | Figure S3A (GFP fusion growth) |
| SBY6313 | *MATa ho::hisG leu2::hisG ura3(ΔSma-Pst) HIS4::LEU2-(NBam) nup60(1-388)-GFP::KanMx*<br>*MATalpha ho::hisG leu2::hisG ura3(ΔSma-Pst) his4-X::LEU2-(NBam)-URA3 nup60(1-388)-GFP::KanMx* | SK1 | Figure S3A (GFP fusion growth) |
| SBY6316 | *MATa ho::hisG leu2::hisG ura3(ΔSma-Pst) HIS4::LEU2-* | SK1 | |

| | | | |
|---|---|---|---|
| | *(NBam) nup60(189-539)-GFP::KanMx*<br>*MATalpha ho::hisG leu2::hisG ura3(ΔSma-Pst) his4-X::LEU2-(NBam)-URA3 nup60(189-539)-GFP::KanMx* | | Figure S3A (GFP fusion growth) |
| SBY6382 | *MATa ho::hisG leu2::hisG ura3(ΔSma-Pst) HIS4::LEU2-(NBam) nup60(Δ189-388)-GFP::KanMx*<br>*MATalpha ho::hisG leu2::hisG ura3(ΔSma-Pst) his4-X::LEU2-(NBam)-URA3 nup60(Δ189-388)-GFP::KanMx* | SK1 | Figure S3A (GFP fusion growth) |
| SBY6320 | *MATa ho::hisG leu2::hisG ura3(ΔSma-Pst) HIS4::LEU2-(NBam) ndj1::KanMx NUP60-GFP::KanMx*<br>*MATalpha ho::hisG leu2::hisG ura3(ΔSma-Pst) his4-X::LEU2-(NBam)-URA3 ndj1::KanMx NUP60-GFP::KanMx* | SK1 | Figure S3B (GFP fusion sporulation) |
| SBY6323 | *MATa ho::hisG leu2::hisG ura3(ΔSma-Pst) HIS4::LEU2-(NBam) ndj1::KanMx nup60(Δ1-539)-GFP::KanMx*<br>*MATalpha ho::hisG leu2::hisG ura3(ΔSma-Pst) his4-X::LEU2-(NBam)-URA3 ndj1::KanMx nup60(Δ1-539)-GFP::KanMx* | SK1 | Figure S3B (GFP fusion sporulation) |
| SBY6326 | *MATa ho::hisG leu2::hisG ura3(ΔSma-Pst) HIS4::LEU2-(NBam) ndj1::KanMx nup60(189-388)-GFP::KanMx*<br>*MATalpha ho::hisG leu2::hisG ura3(ΔSma-Pst) his4-X::LEU2-(NBam)-URA3 ndj1::KanMx nup60(189-388)-GFP::KanMx* | SK1 | Figure S3B (GFP fusion sporulation) |
| SBY6329 | *MATa ho::hisG leu2::hisG ura3(ΔSma-Pst) HIS4::LEU2-(NBam) ndj1::KanMx nup60(1-388)-GFP::KanMx*<br>*MATalpha ho::hisG leu2::hisG ura3(ΔSma-Pst) his4-X::LEU2-(NBam)-URA3 ndj1::KanMx nup60(1-388)-GFP::KanMx* | SK1 | Figure S3B (GFP fusion sporulation) |
| SBY6332 | *MATa ho::hisG leu2::hisG ura3(ΔSma-Pst) HIS4::LEU2-(NBam) ndj1::KanMx nup60(189-539)-GFP::KanMx* | SK1 | Figure S3B (GFP fusion sporulation) |

| | | | | |
|---|---|---|---|---|
| | *MATalpha ho::hisG leu2::hisG ura3(ΔSma-Pst) his4-X::LEU2-(NBam)-URA3 ndj1::KanMx nup60(189-539)-GFP::KanMx* | | | |
| SBY6335 | *MATa ho::hisG leu2::hisG ura3(ΔSma-Pst) HIS4::LEU2-(NBam) ndj1::KanMx nup60(Δ189-388)-GFP::KanMx* | SK1 | Figure S3B (GFP fusion sporulation) | |
| | *MATalpha ho::hisG leu2::hisG ura3(ΔSma-Pst) his4-X::LEU2-(NBam)-URA3 ndj1::KanMx nup60(Δ189-388)-GFP::KanMx* | | | |
| SBY3945 | *MATa ho::hisG leu2::hisG ura3(ΔSma-Pst) HIS4::LEU2-(NBam) nup2::KanMx* | SK1 | Figure 5 (Meiotic time courses) | (Chu *et al.* 2017) |
| | *MATalpha ho::hisG leu2::hisG ura3(ΔSma-Pst) his4-X::LEU2-(NBam)-URA3 nup2::KanMx* | | | |
| SBY6432 | *MATa ho::hisG leu2::hisG ura3(ΔSma-Pst) HIS4::LEU2-(NBam) nup2::KanMx nup60(189-388)* | SK1 | Figure 5 (Meiotic time courses) | |
| | *MATalpha ho::hisG leu2::hisG ura3(ΔSma-Pst) his4-X::LEU2-(NBam)-URA3 nup2::KanMx nup60(189-388)* | | | |
| SBY3983 | *MATa ho::hisG leu2::hisG ura3(ΔSma-Pst) HIS4::LEU2-(NBam) nup2::KanMx ndj1::KanMx* | SK1 | Figure 5 (Meiotic time courses) | (Chu *et al.* 2017) |
| | *MATalpha ho::hisG leu2::hisG ura3(ΔSma-Pst) his4-X::LEU2-(NBam)-URA3 nup2::KanMx ndj1::KanMx* | | | |
| SBY6296 | *MATa ho::hisG leu2::hisG ura3(ΔSma-Pst) HIS4::LEU2-(NBam) ndj1::KanMx nup60(189-388)* | SK1 | Figure 5 (Meiotic time courses) | |
| | *MATalpha ho::hisG leu2::hisG ura3(ΔSma-Pst) his4-X::LEU2-(NBam)-URA3 ndj1::KanMx nup60(189-388)* | | | |
| SBY6254 | *MATa ho::hisG leu2::hisG ura3(ΔSma-Pst) HIS4::LEU2-(NBam) NUP60-GFP::KanMx nup2(51-175)-mCherry::Nat ndt80::Hph* | SK1 | Figures 4C and 6 (NUP60-GFP localization and spreads) | |
| | *MATalpha ho::hisG leu2::hisG ura3(ΔSma-Pst) his4-X::LEU2-(NBam)-URA3 NUP60-GFP::KanMx nup2(51-175)-mCherry::Nat ndt80::Hph* | | | |

| | | | |
|---|---|---|---|
| SBY6338 | *MATa ho::hisG leu2::hisG ura3(ΔSma-Pst) HIS4::LEU2-(NBam) nup60(Δ1-539)-GFP::KanMx nup2(51-175)-mCherry::Nat ndt80::Hph*<br>*MATalpha ho::hisG leu2::hisG ura3(ΔSma-Pst) his4-X::LEU2-(NBam)-URA3 nup60(Δ1-539)-GFP::KanMx nup2(51-175)-mCherry::Nat ndt80::Hph* | SK1 | Figures 4C and 6 (NUP60-GFP localization and spreads) |
| SBY6344 | *MATa ho::hisG leu2::hisG ura3(ΔSma-Pst) HIS4::LEU2-(NBam) nup60(1-388)-GFP::KanMx nup2(51-175)-mCherry::Nat ndt80::Hph*<br>*MATalpha ho::hisG leu2::hisG ura3(ΔSma-Pst) his4-X::LEU2-(NBam)-URA3 nup60(1-388)-GFP::KanMx nup2(51-175)-mCherry::Nat ndt80::Hph* | SK1 | Figures 4C and 6 (NUP60-GFP localization and spreads) |
| SBY6341 | *MATa ho::hisG leu2::hisG ura3(ΔSma-Pst) HIS4::LEU2-(NBam) nup60(189-388)-GFP::KanMx nup2(51-175)-mCherry::Nat ndt80::Hph*<br>*MATalpha ho::hisG leu2::hisG ura3(ΔSma-Pst) his4-X::LEU2-(NBam)-URA3 nup60(189-388)-GFP::KanMx nup2(51-175)-mCherry::Nat ndt80::Hph* | SK1 | Figures 4C and 6 (NUP60-GFP localization and spreads) |
| SBY6347 | *MATa ho::hisG leu2::hisG ura3(ΔSma-Pst) HIS4::LEU2-(NBam) nup60(189-539)-GFP::KanMx nup2(51-175)-mCherry::Nat ndt80::Hph*<br>*MATalpha ho::hisG leu2::hisG ura3(ΔSma-Pst) his4-X::LEU2-(NBam)-URA3 nup60(189-539)-GFP::KanMx nup2(51-175)-mCherry::Nat ndt80::Hph* | SK1 | Figures 4C and 6 (NUP60-GFP localization and spreads) |
| SBY6350 | *MATa ho::hisG leu2::hisG ura3(ΔSma-Pst) HIS4::LEU2-(NBam) nup60(Δ189-388)-GFP::KanMx nup2(51-175)-mCherry::Nat ndt80::Hph*<br>*MATalpha ho::hisG leu2::hisG ura3(ΔSma-Pst) his4-X::LEU2-(NBam)-URA3 nup60(Δ189-388)-GFP::KanMx nup2(51-175)-mCherry::Nat ndt80::Hph* | SK1 | Figure 4C (NUP60-GFP localization) |

| | | | |
|---|---|---|---|
| SBY6365 | *MATa ho::hisG leu2::hisG ura3(ΔSma-Pst) HIS4::LEU2-(NBam) ndj1::KanMx nup2::Nat NUP60-GFP::KanMx*<br>*MATalpha ho::hisG leu2::hisG ura3(ΔSma-Pst) his4-X::LEU2-(NBam)-URA3 ndj1::KanMx nup2::Nat NUP60-GFP::KanMx* | SK1 | Table 3 (Suppression of nup2 ndj1) |
| SBY6371 | *MATa ho::hisG leu2::hisG ura3(ΔSma-Pst) HIS4::LEU2-(NBam) ndj1::KanMx nup2::Nat nup60(1-388)-GFP::KanMx*<br>*MATalpha ho::hisG leu2::hisG ura3(ΔSma-Pst) his4-X::LEU2-(NBam)-URA3 ndj1::KanMx nup2::Nat nup60(1-388)-GFP::KanMx* | SK1 | Table 3 (Suppression of nup2 ndj1) |
| SBY6356 | *MATa ho::hisG leu2::hisG ura3(ΔSma-Pst) HIS4::LEU2-(NBam) ndj1::KanMx nup2::Nat nup60(1-388)*<br>*MATalpha ho::hisG leu2::hisG ura3(ΔSma-Pst) his4-X::LEU2-(NBam)-URA3 ndj1::KanMx nup2::Nat nup60(1-388)* | SK1 | Table 3 (Suppression of nup2 ndj1) |
| SBY6436 | *MATa ho::hisG leu2::hisG ura3(ΔSma-Pst) HIS4::LEU2-(NBam) ndj1::KanMx nup2::Nat NUP60-mCherry::KanMx*<br>*MATalpha ho::hisG leu2::hisG ura3(ΔSma-Pst) his4-X::LEU2-(NBam)-URA3 ndj1::KanMx nup2::Nat NUP60-mCherry::KanMx* | SK1 | Table 3 (Suppression of nup2 ndj1) |
| SBY6439 | *MATa ho::hisG leu2::hisG ura3(ΔSma-Pst) HIS4::LEU2-(NBam) ndj1::KanMx nup2::Nat NUP60-13myc::KanMx*<br>*MATalpha ho::hisG leu2::hisG ura3(ΔSma-Pst) his4-X::LEU2-(NBam)-URA3 ndj1::KanMx nup2::Nat NUP60-13myc::KanMx* | SK1 | Table 3 (Suppression of nup2 ndj1) |
| SBY3982 x SBY6364 | *MATa ho::hisG leu2::hisG ura3(ΔSma-Pst) HIS4::LEU2-(NBam) ndj1::KanMx nup2::Nat*<br>*MATalpha ho::hisG leu2::hisG ura3(ΔSma-Pst) his4-X::LEU2-(NBam)-URA3 ndj1::KanMx nup2::Nat NUP60-GFP::KanMxB* | SK1 | Table 3 (Suppression of nup2 ndj1) |
| SBY3982 x SBY6435 | *MATa ho::hisG leu2::hisG ura3(ΔSma-Pst) HIS4::LEU2-(NBam) ndj1::KanMx nup2::Nat* | SK1 | Table 3 (Suppression of nup2 ndj1) |

| | | | |
|---|---|---|---|
| | *MATalpha ho::hisG leu2::hisG ura3(ΔSma-Pst) his4-X::LEU2-(NBam)-URA3 ndj1::KanMx nup2::Nat NUP60-mCherry::KanMx* | | |
| SBY3982 x SBY6370 | *MATa ho::hisG leu2::hisG ura3(ΔSma-Pst) HIS4::LEU2-(NBam) ndj1::KanMx nup2::Nat* | SK1 | Table 3 (Suppression of nup2 ndj1) |
| | *MATalpha ho::hisG leu2::hisG ura3(ΔSma-Pst) his4-X::LEU2-(NBam)-URA3 ndj1::KanMx nup2::Nat nup60(1-388)-GFP::KanMx* | | |
| SBY3982 x SBY6355 | *MATa ho::hisG leu2::hisG ura3(ΔSma-Pst) HIS4::LEU2-(NBam) ndj1::KanMx nup2::Nat* | SK1 | Table 3 (Suppression of nup2 ndj1) |
| | *MATalpha ho::hisG leu2::hisG ura3(ΔSma-Pst) his4-X::LEU2-(NBam)-URA3 ndj1::KanMx nup2::Nat nup60(1-388)* | | |
| SBY6305 | *MATa ho::hisG leu2::hisG ura3(ΔSma-Pst) HIS4::LEU2-(NBam) NUP60-GFP::KanMx* | SK1 | Figure 7 (Nup60-GFP binding with and without Nup2) |
| | *MATalpha ho::hisG leu2::hisG ura3(ΔSma-Pst) his4-X::LEU2-(NBam)-URA3 NUP60-GFP::KanMx* | | |
| SBY6487 | *MATa ho::hisG leu2::hisG ura3(ΔSma-Pst) HIS4::LEU2-(NBam) NUP60-GFP::KanMx nup2::NatMx* | SK1 | Figure 7 (Nup60-GFP binding with and without Nup2) |
| | *MATalpha ho::hisG leu2::hisG ura3(ΔSma-Pst) his4-X::LEU2-(NBam)-URA3 NUP60-GFP::KanMx nup2::NatMx* | | |
| SBY6311 | *MATa ho::hisG leu2::hisG ura3(ΔSma-Pst) HIS4::LEU2-(NBam) nup60(189-388)-GFP::KanMx* | SK1 | Figure 7 (Nup60-GFP binding with and without Nup2) |
| | *MATalpha ho::hisG leu2::hisG ura3(ΔSma-Pst) his4-X::LEU2-(NBam)-URA3 nup60(189-388)-GFP::KanMx* | | |
| SBY6442 | *MATa ho::hisG leu2::hisG ura3(ΔSma-Pst) HIS4::LEU2-(NBam) csm4::Hph* | SK1 | Table S2 (nup60 epistasis) |
| | *MATalpha ho::hisG leu2::hisG ura3(ΔSma-Pst) his4-X::LEU2-(NBam)-URA3 csm4::Hph* | | |
| SBY6448 | *MATa ho::hisG leu2::hisG ura3(ΔSma-Pst) HIS4::LEU2-(NBam) nup60(189-388) csm4::Hph* | SK1 | Table S2(nup60 epistasis) |

| | | | |
|---|---|---|---|
| | *MATalpha ho::hisG leu2::hisG ura3(ΔSma-Pst) his4-X::LEU2-(NBam)-URA3 nup60(189-388) csm4::Hph* | | |
| SBY6445 | *MATa ho::hisG leu2::hisG ura3(ΔSma-Pst) HIS4::LEU2-(NBam) nup60Δ csm4::Hph* | SK1 | Table S2 (nup60 epistasis) |
| | *MATalpha ho::hisG leu2::hisG ura3(ΔSma-Pst) his4-X::LEU2-(NBam)-URA3 nup60Δ csm4::Hph* | | |
| SBY6451 | *MATa ho::hisG leu2::hisG ura3(ΔSma-Pst) HIS4::LEU2-(NBam) nup2::NatMx csm4::Hph* | SK1 | Table S2 (nup60 epistasis) |
| | *MATalpha ho::hisG leu2::hisG ura3(ΔSma-Pst) his4-X::LEU2-(NBam)-URA3 nup2::NatMx csm4::Hph* | | |
| SBY6454 | *MATa ho::hisG leu2::hisG ura3(ΔSma-Pst) HIS4::LEU2-(NBam) nup60(189-388) nup2::KanMx* | SK1 | Table S2 (nup60 epistasis) |
| | *MATalpha ho::hisG leu2::hisG ura3(ΔSma-Pst) his4-X::LEU2-(NBam)-URA3 nup60(189-388) nup2::KanMx* | | |
| SBY6353 | *MATa ho::hisG leu2::hisG ura3(ΔSma-Pst) HIS4::LEU2-(NBam) nup60(189-388) nup2::KanMx ndj1::KanMx* | SK1 | Table S2 (nup60 epistasis) |
| | *MATalpha ho::hisG leu2::hisG ura3(ΔSma-Pst) his4-X::LEU2-(NBam)-URA3 nup60(189-388) nup2::KanMx ndj1::KanMx* | | |
| SBY6457 | *MATa ho::hisG leu2::hisG ura3(ΔSma-Pst) HIS4::LEU2-(NBam) 3HA-NUP60* | SK1 | Figure S5A |
| | *MATalpha ho::hisG leu2::hisG ura3(ΔSma-Pst) his4-X::LEU2-(NBam)-URA3 3HA-NUP60* | | |
| SBY6460 | *MATa ho::hisG leu2::hisG ura3(ΔSma-Pst) HIS4::LEU2-(NBam) 3HA-NUP60 ndj1::Hph* | SK1 | Figure S5A |
| | *MATalpha ho::hisG leu2::hisG ura3(ΔSma-Pst) his4-X::LEU2-(NBam)-URA3 3HA-NUP60 ndj1::Hph* | | |
| SBY6463 | *MATa ho::hisG leu2::hisG ura3(ΔSma-Pst) HIS4::LEU2-(NBam) 3HA-NUP60 nup2::NatMx* | SK1 | Figure S5A |
| | *MATalpha ho::hisG leu2::hisG ura3(ΔSma-Pst) his4-X::LEU2-* | | |

| | | | |
|---|---|---|---|
| | *(NBam)-URA3 3HA-NUP60 nup2::NatMx* | | |
| SBY6466 | *MATa ho::hisG leu2::hisG ura3(ΔSma-Pst) HIS4::LEU2-(NBam) 3HA-NUP60 ndj1::Hph nup2::NatMx*<br>*MATalpha ho::hisG leu2::hisG ura3(ΔSma-Pst) his4-X::LEU2-(NBam)-URA3 3HA-NUP60 ndj1::Hph nup2::NatMx* | SK1 | Figure S5A |
| SBY6469 | *MATa ho::hisG leu2::hisG ura3(ΔSma-Pst) HIS4::LEU2-(NBam) NUP60-3HA(Δ1F)::KanMx*<br>*MATalpha ho::hisG leu2::hisG ura3(ΔSma-Pst) his4-X::LEU2-(NBam)-URA3 NUP60-3HA(Δ1F)::KanMx* | SK1 | Figure S5B |
| SBY6472 | *MATa ho::hisG leu2::hisG ura3(ΔSma-Pst) HIS4::LEU2-(NBam) NUP60-3HA(Δ1F)::KanMx ndj1::Hph*<br>*MATalpha ho::hisG leu2::hisG ura3(ΔSma-Pst) his4-X::LEU2-(NBam)-URA3 NUP60-3HA(Δ1F)::KanMx ndj1::Hph* | SK1 | Figure S5B |
| SBY6475 | *MATa ho::hisG leu2::hisG ura3(ΔSma-Pst) HIS4::LEU2-(NBam) NUP60-3HA(Δ1F)::KanMx nup2::NatMx*<br>*MATalpha ho::hisG leu2::hisG ura3(ΔSma-Pst) his4-X::LEU2-(NBam)-URA3 NUP60-3HA(Δ1F)::KanMx nup2::NatMx* | SK1 | Figure S5B |
| SBY6478 | *MATa ho::hisG leu2::hisG ura3(ΔSma-Pst) HIS4::LEU2-(NBam) NUP60-3HA(Δ1F)::KanMx nup2::NatMx ndj1::Hph*<br>*MATalpha ho::hisG leu2::hisG ura3(ΔSma-Pst) his4-X::LEU2-(NBam)-URA3 NUP60-3HA(Δ1F)::KanMx nup2::NatMx ndj1::Hph* | SK1 | Figure S5B |
| SBY6481 | *MATa ho::hisG leu2::hisG ura3(ΔSma-Pst) HIS4::LEU2-(NBam) NUP60-mCherry::KanMx nup2(51-175)-GFP::CaURA3 ndt80::Hph*<br>*MATalpha ho::hisG leu2::hisG ura3(ΔSma-Pst) his4-X::LEU2-(NBam)-URA3 NUP60-* | SK1 | Figure S6 (Nup60-mCherry spreads) |

| | | | |
|---|---|---|---|
| | *mCherry::KanMx nup2(51-175)-GFP::CaURA3 ndt80::Hph* | | |
| SBY6484 | *MATa ho::hisG leu2::hisG ura3(ΔSma-Pst) HIS4::LEU2-(NBam) NUP60-3HA::KanMx nup2(51-175)-GFP::CaURA3 ndt80::Hph*<br>*MATalpha ho::hisG leu2::hisG ura3(ΔSma-Pst) his4-X::LEU2-(NBam)-URA3 NUP60-3HA::KanMx nup2(51-175)-GFP::CaURA3 ndt80::Hph* | SK1 | Figure S6 (Nup60-3HA spreads) |

**Table S2.** Plasmids used in this study

| Plasmid name | Description | Source/ Reference |
|---|---|---|
| pCR-BluntII-TOPO | cloning vector | Thermo Fisher/Invitrogen |
| pSB740 | MAR-GFP::URA3 subcloned into pCR-Blunt II-TOPO | |
| pGBKT7 | Gal4 DBD vector | Clontech |
| pACT2-2 | Gal4 AD vector | (Arora *et al.* 2004) |
| pSB655 | SK1 Nup2(51-175) coding sequence subcloned into pGBKT7 | |
| pSB745 | Nup2(51-175)$^{F100S}$ coding sequence subcloned into pGBKT7 | |
| pSB746 | Nup2(51-175)$^{K101E}$ coding sequence subcloned into pGBKT7 | |
| pSB747 | Nup2(51-175)$^{R117W}$ coding sequence subcloned into pGBKT7 | |
| pSB748 | Nup2(51-175)$^{L98P}$ coding sequence subcloned into pGBKT7 | |
| pSB749 | Nup2(51-175)$^{A102P}$ coding sequence subcloned into pGBKT7 | |
| pSB750 | Nup2(51-175)$^{L119P}$ coding sequence subcloned into pGBKT7 | |
| pSB751 | Nup2(51-175)$^{Y123C}$ coding sequence subcloned into pGBKT7 | |
| pSB752 | Nup2(51-175)$^{D115V}$ coding sequence subcloned into pGBKT7 | |
| pSB753 | Nup2(51-175)$^{L93P}$ coding sequence subcloned into pGBKT7 | |
| pSB754 | Nup2(51-175)$^{L96S}$ coding sequence subcloned into pGBKT7 | |
| pSB755 | Nup2(51-175)$^{N97S}$ coding sequence subcloned into pGBKT7 | |
| pSB756 | Nup2(51-175)$^{V108D}$ coding sequence subcloned into pGBKT7 | |
| pSB757 | Nup2(51-175)$^{F120Y}$ coding sequence subcloned into pGBKT7 | |
| pSB758 | Nup2(51-175)$^{Y126N}$ coding sequence subcloned into pGBKT7 | |
| pSB759 | Nup2(51-175)$^{L116S}$ coding sequence subcloned into pGBKT7 | |
| pSB743 | Nup2(71-130) coding sequence subcloned into pGBKT7 | |
| pSB744 | Nup2(89-130) coding sequence subcloned into pGBKT7 | |
| pSB741 | SK1 Nup60 coding sequence subcloned into pACT2-2 | |

| | |
|---|---|
| pSB742 | SK1 Nup60(188-388) coding sequence subcloned into pACT2-2 |

**Table S3.** Primers for PCR mutagenesis and Y2H plasmid construction

| Primer name | Sequence | Description |
|---|---|---|
| oSB1274 | GGGTTCGACTCCCCGTATC | forward primer for amplifying the full MAR-GFP::URA3 fragment<br>MAR mutagenesis: forward primer for amplifying the 5' fragment containing the MAR |
| oSB1774 | TGCGCGATGTTTTAACGAAG | reverse primer for amplifying the full MAR-GFP::URA3 fragment<br>MAR mutagenesis: reverse primer for amplifying the 3' fragment containing GFP-URA3 |
| oSB1918 | TAAACCAGCACCGTCACCC | MAR mutagenesis: reverse primer for amplifying the 5' fragment containing MAR |
| oSB1919 | GGGTGACGGTGCTGGTTTA | MAR mutagenesis: forward primer for amplifying the 3' fragment containing GFP-URA3 |
| oSB1616 | ACTGGAATTCATGAAACCTTTTGGTTCTGCAA | Y2H: forward primer for insertion of Nup2(51-175) coding sequence into pGBKT7 |
| oSB1617 | ACTGGGATCCTTACTTGGGTCCCTCCACCTTAACC | Y2H: reverse primer for insertion of Nup2(51-175) coding sequence into pGBKT7 |
| oSB1920 | ACTGCAACTAGTATGCATCGTAAATCATTGAGGAGGGCTA G | Y2H: forward primer for insertion of Nup60 coding sequence into pACT2-2 |
| oSB1921 | TAAGCACCATGGCAAAGGTATATAGGGACTTGAAAGCCTC AAC | Y2H: reverse primer for insertion of Nup60 coding sequence into pACT2-2 |
| oSB1922 | ACTGCAACTAGTATGCCGACCTTCAACCCAAAATATGATAC TTCAAATG | Y2H: forward primer for insertion of Nup60(188-388) |

| | | |
|---|---|---|
| | | coding sequence into pACT2-2 |
| oSB1923 | TAAGCACCATGGCACCATCCTTCTTTTCAGGTGAGGTTTCAG | Y2H: reverse primer for insertion of Nup60(188-388) coding sequence into pACT2-2 |
| oSB1924 | ACTGGAATTCATGAACCGGGCGGACGGCACTG | Y2H: forward primer for insertion of Nup2(71-130) coding sequence into pGBKT7 |
| oSB1925 | ACTGGAATTCATGAGCAATTCCAGACTAAAAGCATTGAACC | Y2H: forward primer for insertion of Nup2(89-130) coding sequence into pGBKT7 |
| oSB1926 | ACTGGGATCCTTAGATATTCTTTATGTATAATTCGTACCTGGTG | Y2H: reverse primer for insertion of Nup2(71-130) and Nup2(89-130) coding sequences into pGBKT7 |

### Supplemental methods

#### Western blot

Yeast extracts were prepared by alkaline lysis (von der Haar 2007) and resolved on a 4-15% Mini-PROTEAN TGX gel (BioRad, 4561085). Gel electrophoresis and wet transfer to PVDF were carried out according to standard procedures (Bolt and Mahoney 1997). Western blot detection was performed using ECL reagents and the manufacturer's instructions for PBS-based buffers (Amersham, RPN2108). Membranes were imaged with an ImageQuant LAS4000.

#### Antibodies

Primary antibodies used were mouse monoclonal antibody to HA (Santa Cruz Biotechnology SC-7392; RRID:AB_627809), rabbit polyclonal antibody to β-tubulin (Abcam AB15568; RRID:AB_2210952), chicken polyclonal antibody to GFP (Novus NB100-1614; RRID:AB_10001164), and rabbit polyclonal antibody to mCherry (Novus NBP2-25157; RRID:AB_2753204). The secondary antibodies used were goat anti-chicken 488 (Thermofisher A11039; RRID:AB_2534096), goat anti-rabbit 594 (ThermofisherA11012; RRID:AB_2534079), donkey anti-mouse 488 (Invitrogen A32766; RRID:AB_2762823), and the anti-rabbit and anti-mouse HRP-conjugated secondary antibodies provided in the ECL kit. All antibodies were used at a 1:1000 dilution.

### Supplemental references

Arora C., K. Kee, S. Maleki, and S. Keeney, 2004 Antiviral protein Ski8 is a direct partner of Spo11 in meiotic DNA break formation, independent of its cytoplasmic role in RNA metabolism. Mol. Cell 13: 549–559.

Bolt M. W., and P. A. Mahoney, 1997 High-efficiency blotting of proteins of diverse sizes following sodium dodecyl sulfate-polyacrylamide gel electrophoresis. Anal. Biochem. 247: 185–192.

Chu D. B., T. Gromova, T. A. C. Newman, and S. M. Burgess, 2017 The Nucleoporin Nup2 Contains a Meiotic-Autonomous Region that Promotes the Dynamic Chromosome Events of Meiosis. Genetics 206: 1319–1337.

Haar T. von der, 2007 Optimized protein extraction for quantitative proteomics of yeasts. PLoS One 2: e1078.

Jumper J., R. Evans, A. Pritzel, T. Green, M. Figurnov, *et al.*, 2021 Highly accurate protein structure prediction with AlphaFold. Nature 596: 583–589.

Mariani V., M. Biasini, A. Barbato, and T. Schwede, 2013 lDDT: a local superposition-free score for comparing protein structures and models using distance difference tests. Bioinformatics 29: 2722–2728.

Pettersen E. F., T. D. Goddard, C. C. Huang, E. C. Meng, G. S. Couch, *et al.*, 2021 UCSF ChimeraX: Structure visualization for researchers, educators, and developers. Protein Sci. 30: 70–82.

Saiz-Baggetto S., E. Méndez, I. Quilis, J. C. Igual, and M. C. Bañó, 2017 Chimeric proteins tagged with specific 3xHA cassettes may present instability and functional problems. PLoS One 12: e0183067.
